# Strawberry COP9 signalosome FvCSN5A regulates plant development and fruit ripening by facilitating polyamine oxidase FvPAO5 degradation to control polyamine and H_2_O_2_ homeostasis

**DOI:** 10.1101/2024.07.10.602942

**Authors:** Yun Huang, Jiahui Gao, Guiming Ji, Wenjing Li, Jiaxue Wang, Qinghua Wang, Yuanyue Shen, Jiaxuan Guo, Fan Gao

## Abstract

Polyamines (PAs), mainly including putrescine, spermidine, and spermine, are essential for plant growth and development. However, the post-translational regulation of PA metabolism remains unknown. Here, we report the COP9 signalosome subunit 5A (FvCSN5A)-mediated degradation of the PA oxidase FvPAO5, which catalyzes the conversion of spermidine/spermine to produce H_2_O_2_. FvCSN5A is localized in the cytoplasm and nucleus, ubiquitously expressed in strawberry plants, and rapidly increases during fruit ripening. *FvCSN5A* RNA interference (RNAi) transgenic strawberries exhibit pleiotropic effects on plant development, fertility, and fruit ripening by regulating PA and H_2_O_2_ homeostasis, similar to *FvPAO5* overexpression transgenic lines. FvCSN5A interacts with FvPAO5 in vitro and in vivo. The ubiquitination and degradation of FvPAO5 are impaired in *FvCSN5A* RNAi lines. Additionally, FvCSN5A interacts with cullin 1 (FvCUL1), a core component of the E3 ubiquitin-protein ligase complex. Transient genetic analysis in cultivated strawberry fruits showed that inhibiting *FaPAO5* expression could partially rescue the ripening phenotype of *FaCSN5A* RNAi fruits. Collectively, the CSN5A-CUL1-PAO5 signaling pathway responsible for PA and H_2_O_2_ homeostasis is crucial to strawberry vegetative and reproductive growth and, final to fruit ripening. Our findings may open a promising avenue for improving crop yield and quality by manipulating the CSN5-PAO5 pair.

## Introduction

Polyamines (PAs) are small polycationic molecules, primarily consisting of putrescine (Put), spermidine (Spd), and spermine (Spm). These molecules have been demonstrated to participate in numerous molecular processes, such as serving as a nitrogen storage pool (Siddappa and Marathe, 2020), protecting and stabilizing macromolecules (Napieraj et al., 2023), regulating reactive oxygen species (ROS) homeostasis (Pottosin et al., 2014; Jasso-Robles et al., 2020), and participating in signal transduction (Gao et al., 2021). Consequently, PAs play important roles in various aspects of plant development and growth, including photosynthetic regulation, secondary metabolism, flowering, fruit ripening, and environmental responses (Fortes et al., 2019; Wang et al., 2019; Aloisi et al., 2022; Nandy et al., 2022; Navakoudis and Kotzabasis, 2022; Blázquez, 2024). Although much progress has been made in understanding PA metabolism and function, the regulatory mechanisms governing PA homeostasis at the post-translational level remain elusive.

In general, Put is involved in photosynthetic protection, while Spd/Spm, especially Spm, are involved in regulating ROS (Zhao et al., 2021; Zhong et al., 2023; Blázquez, 2024; Furumoto et al., 2024). In response to developmental and environmental stimuli, PA equilibria are fine-tuned at biosynthetic and degradative levels (Gupta et al., 2016; Wang et al., 2019; Zhan et al., 2023). That is, PA biosynthesis begins with Put, catalyzed by either arginine decarboxylase (ADC) or ornithine decarboxylase (ODC). Then, Put is utilized as a substrate for the successive biosynthesis of Spd and Spm, catalyzed separately by Spd synthase (SPDS) and Spm synthase (SPMS). This process requires the addition of an aminopropyl group derived from decarboxylated S-adenosyl-L-methionine (dcSAM). The dcSAM is generated from S-adenosyl-L-methionine (SAM), catalyzed by SAM decarboxylase (SAMDC) using SAM as a substrate (Liu et al., 2006; Groppa and Benavides, 2008; Gupta et al., 2016). PA degradation involves copper-containing diamine oxidase (CuAO/DAO)-mediated Put degradation and flavin-containing polyamine oxidase (PAO)-mediated Spd/Spm degradation in a back-conversion (BC-type) or terminal-catabolism (TC-type) manner (Paschalidis et al., 2005; Angelini et al., 2010; Yu et al., 2019). In particular, PA decomposition produces H_2_O_2_, which contributes to fine-tuning plant growth and development in response to environmental cues by PA and ROS homeostasis (Navakoudis and Kotzabasis, 2022; Zhan et al.,

Notably, AtPAO5 has been demonstrated to participate in Arabidopsis xylem differentiation by auxin and cytokinin (Alabdallah et al., 2017). Similarly, OsPAO5 plays an important role in rice mesocotyl and seedling growth, as well as grain weight, grain number, and yield potential through H_2_O_2_ and ethylene (Lv et al., 2021). Additionally, PAO5 acts as a negative regulator during strawberry fruit ripening by the TC-type degradation of Spm/Spd in concomitant with H_2_O_2_ production (Mo et al., 2020). However, the mechanism behind the continuously-decreased protein levels of FaPAO5 during fruit ripening (Mo et al., 2020) remains unclear, this evokes us to speculate that ubiquitination-mediated degradation of PAO5 might be involved.

The cullin-RING ubiquitin ligase (CRL) pathway, a common protein degradation mechanism, is prevalent in all eukaryotes and facilitates multiple cellular processes through the ubiquitin-mediated degradation of targeted proteins via an E1-E2-E3 cascade of the 26S proteasome system (Barry and Früh, 2006; Hotton and Callis, 2008). Specifically, the 76-amino-acid ubiquitin (UB) is covalently attached to E1 (ubiquitin-activating enzyme) via ATP-derived adenylation, then the activated UB is successively transferred to E2 (ubiquitin-conjugating enzyme) and E3 (ubiquitin ligase) complex.

Within the E3 ligase superfamily, CRL activity is regulated by RUB/NEDD8 (related to ubiquitin-like protein) and the COP9 signalosome (CSN; Hotton and Callis, 2008). As a component of the ubiquitin-proteasome system, the CSN regulates CRL assembly via neddylation and deneddylation processes (Stratmann and Gusmaroli, 2012). Notably, the CSN complex was initially identified through genetic screening of light□induced photomorphogenic mutants in Arabidopsis, indicating that CSN plays a critical role in controlling CRL activity and various cellular processes (Serino and Deng, 2003; Qin et al., 2020; Lu et al., 2024).

In the COP9 signalosome complex, CSN1-4, CSN7, and CSN8 contain a PCI (proteasome, COP9, initiation factor 3) domain, while CSN5 and CSN6 contain a domain of MPN (Mov34 and Pad1p N□terminal), among which CSN5 acts as a zinc-ion-activating metalloprotease and has two paralogs, designated CSN5A and CSN5B (Verma et al., 2002; Cope and Deshaies, 2003; Wei and Deng, 2003; Gusmaroli et al., 2004; Jin et al., 2014). Notably, CSN5A plays an important role in abscisic acid (ABA) signal transduction by regulating the stability of the bZIP transcription factor ABI5 (Jin et al., 2014). The MPN domain is related to both CSN deneddylation and 26S-proteasome deubiquitination, while the PCI domain is responsible for subunit interaction (Huang et al., 2005). The al., 2014; Reitsma et al., 2017; Liu et al., 2018; Mayor-Ruiz et al., 2019). For example, CSN regulates the NEDD8-modification of CUL-based E3 ligases in SCF^TIR1^, SCF^SLY^, and SCF^COI1^ complexes, facilitating the deneddylation of these CRLs in vivo (Qin et al., 2020). The E3 ubiquitin ligase SCF^TIR1^ directly interacts with the CSN, which is essential for the efficient degradation of its substrates in Arabidopsis (Schwechheimer et al., 2001). A more recent report indicated that the CSN-mediated deneddylation of CUL1 is necessary for the proper assembly of SCF^EBF1^ in Arabidopsis (Dong et al., 2024). CSN5 is localized in the nucleus and cytosol, and can cleave the NEDD8-CUL1 conjugate responsible for deneddylation and protein degradation (Kwok et al., 1998; Lyapina et al., 2001; Schwechheimer et al., 2001). The CSN-mediated deneddylation not only prevents auto-ubiquitination of CRL components, but also facilitates the recycling of the cullin-RING heterodimer for numerous F-box/substrate interactions (Fischer et al., 2011). Intriguingly, two more recent studies find that CSN5 inhibits autophagy by regulating the ubiquitination of Atg6 (autophagy-related) and TOR (target of rapamycin), thereby mediating the pathogenicity of *Magnaporthe oryzae* (Shen et al., 2024). Under stress conditions, CSN5A regulates the stability of chloroplast proteins in tomato (*Solanum lycopersicum*) by degrading SlPsbS (S subunit of PSII) in the cytosol (Lu et al., 2024). Collectively, the central role of CSN5 within the COP9 signalosome involves a wide range of cellular and biological processes, including light signaling, hormone signaling, and environmental responses in plants (Qin et al., 2020). However, whether the COP9 signalosome participates in PA metabolism remains unknown.

To explore the potential mechanism underlying the continued decline of PAO5 during strawberry fruit ripening (Mo et al., 2020), we initially conducted a yeast two-hybrid library screening to identify the interaction protein of PAO5. Following confirmation of the interaction, FvCSN5A was identified through localization, spatiotemporal expression, and RNAi-stable transgenic analyses. Subsequently, the potential relationship between FvCSN5A and FvPAO5 in terms of ubiquitination degradation was explored. Finally, the results obtained from diploid strawberries during fruit ripening were confirmed in octaploid cultivated strawberries. Based on the data available and our current findings, we propose a post-translational regulatory model for PA and H_2_O_2_ homeostasis, controlled by the FvCSN5A-FvCUL1-FvPAO5 pathway. This provides novel insights into the integration of the COP9 signalosome into PA metabolism, opening an attractive avenue for agricultural practices aimed at improving crop yield and quality through manipulation of the CSN5-PAO5 pair.

## Results

### Identification of FvCSN5A as FvPAO5 interacting protein

Given the continual decrease in FaPAO5 protein levels during strawberry fruit ripening (Mo et al., 2020), it is potential to FaPAO5 involved in protein degradation. To investigate this hypothesis, we used diploid strawberries (*Fragaria vesca*, Ruegen) and performed a yeast two-hybrid (Y2H) screen using FvPAO5 as a bait. As a result, we identified an interacting protein of FvPAO5, designated FvCSN5A, which is homologous to the Arabidopsis CSN5A protein (Fig. 1A). We subsequently validated the physical interaction between FvPAO5 and FvCSN5A using a variety of complementary experimental approaches. In the GST pull-down assay conducted in vitro, the FvPAO5-GST fusion protein was precipitated by FvCSN5A-His (Fig. 1B). Furthermore, the firefly luciferase complementation (FLC) assay revealed a strong fluorescent signal upon co-transformation of the FvPAO5-NLUC and FvCSN5A-CLUC constructs in tobacco leaves, while no such signal was observed in the control (Fig. 1C). To corroborate the interaction in vivo, we performed co-immunoprecipitation (CoIP) experiments by co-expressing MYC-tagged FvCSN5A with either FvPAO5-GFP or GFP alone in *Nicotiana benthamiana* epidermal cells via Agroinfiltration. The results demonstrated that MYC-FvCSN5A could specifically associate with FvPAO5-GFP, but not with GFP alone, indicating that FvPAO5 interacts with FvCSN5A (Fig. 1D).

**Figure 1.**
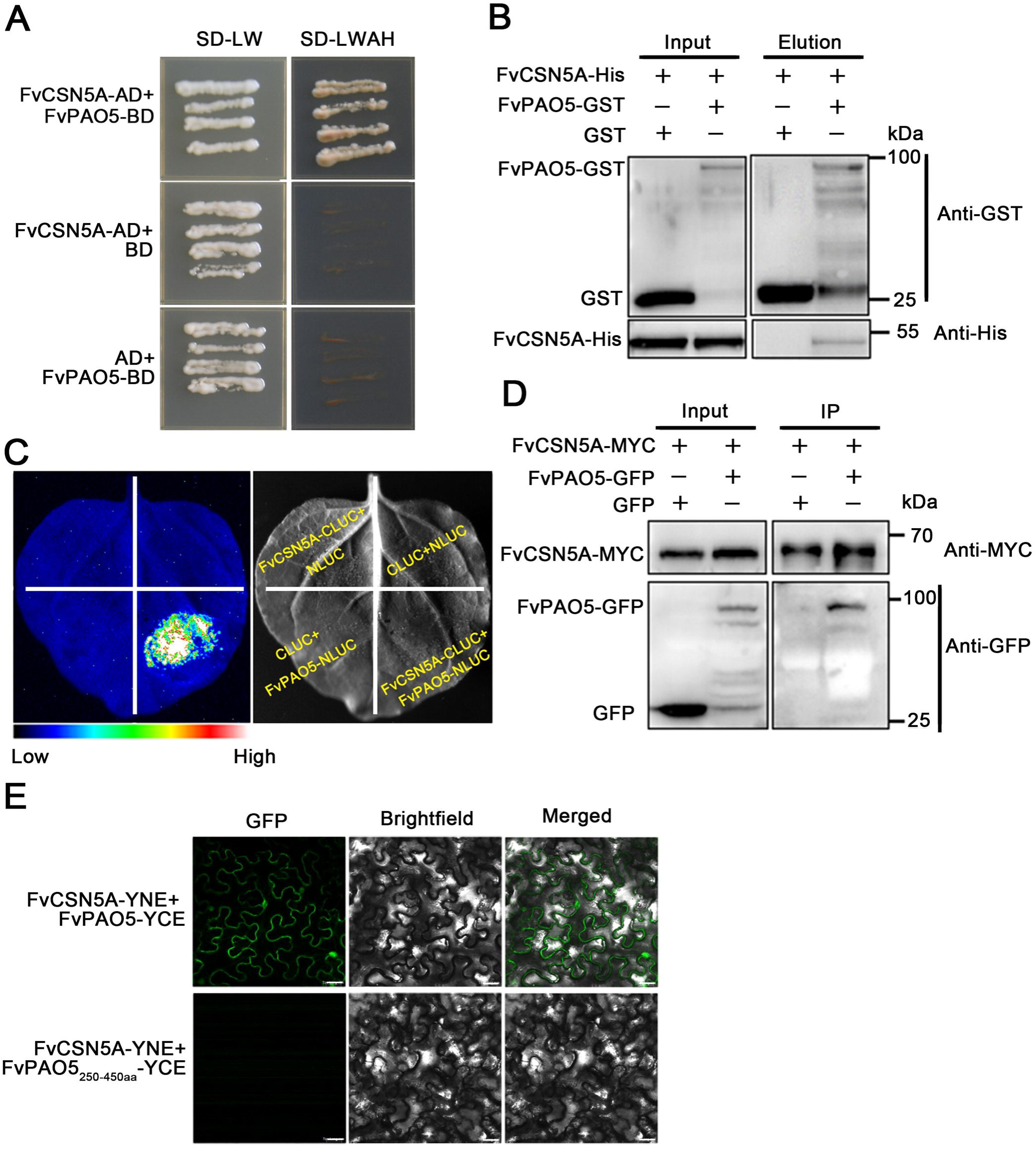
FvCSN5A interacted with FvPAO5 *in vitro* and *in vivo*. **(A)** FvCSN5A interacted with FvPAO5 in yeast two-hybrid assays. SD-LW: synthetic dropout medium without Leu and Trp; SD-LWHA: synthetic dropout medium without Leu, Trp, His, and Ade. (**B)** FvCSN5A interacted with FvPAO5 in *vitro* pull-down assays. Recombinant GST or FvPAO5-GST bounded to glutathione sepharose beads was incubated with recombinant FvCSN5A-His protein. Input and elution proteins immunoblotted with anti-His and anti-GST antibodies, respectively. (**C)** FvCSN5A interacted with FvPAO5 in firefly luciferase complementation (FLC) assays. The CDS of *FvCSN5A* was constructed into the pCAMBIA1300-CLUC vector, and the CDS of *FvPAO5* was constructed into the pCAMBIA1300-NLUC vector. (**D)** FvCSN5A interacted with FvPAO5 in *vivo* Co-Immunoprecipitation (Co-IP) assays. FvCSN5A-MYC protein bounded to MYC beads was incubated with GFP or FvPAO5-GFP and immunoblotted with anti-GFP antibody and anti-MYC antibody, respectively. (**E)** FvCSN5A interacted with FvPAO5 in bimolecular fluorescence complementation (BiFC) assays. FvCSN5A was fused to the N terminus of YFP and FvPAO5 to the C terminus of YFP. FvPAO5_250-450aa_ is a truncated form of FvPAO5 protein that acts as a negative control. The encoding constructs were co-infiltrated into *N. benthamiana* leaves. Bars, 20 μm.

Finally, we employed a bimolecular fluorescence complementation (BiFC) assay to further validate the FvCSN5A-FvPAO5 interaction in plant cells. The FvCSN5A-YNE (N-terminal of YFP) and FvPAO5-YCE (C-terminal of YFP) constructs were delivered into *N. benthamiana* leaf cells, and distinct fluorescence was observed in the cytoplasm and nucleus, providing additional evidence for the interaction between FvPAO5 and FvCSN5A (Fig. 1E). Collectively, these complementary experimental results provide consolidated evidence that FvPAO5 interacts with FvCSN5A both in vitro and in vivo.

### Expression pattern and subcellular localization of FvCSN5A

To explore the expression pattern of *FvCSN5A*, we tracked its transcriptional level in different organs of strawberry plants and across five developmental stages of strawberry fruit using RT-qPCR. The expression level of *FvCSN5A* was highest in roots, followed by red seeds, stems, and leaves, while it was lowest in flowers and fruits (Fig. 2A). The *FvCSN5A* transcript level increased gradually from the white (Wt) to partial red (PR) stage and reached a peak during the full red (FR) stage (Fig. 2B). In contrast, the expression level of *FvPAO5* decreased rapidly from the small green (SG) to FR stage (Mo et al., 2020). These results suggest that FvCSN5A may exert an effect on the fruit ripening process, potentially related to the decreased FvPAO5 levels.

**Figure 2.**
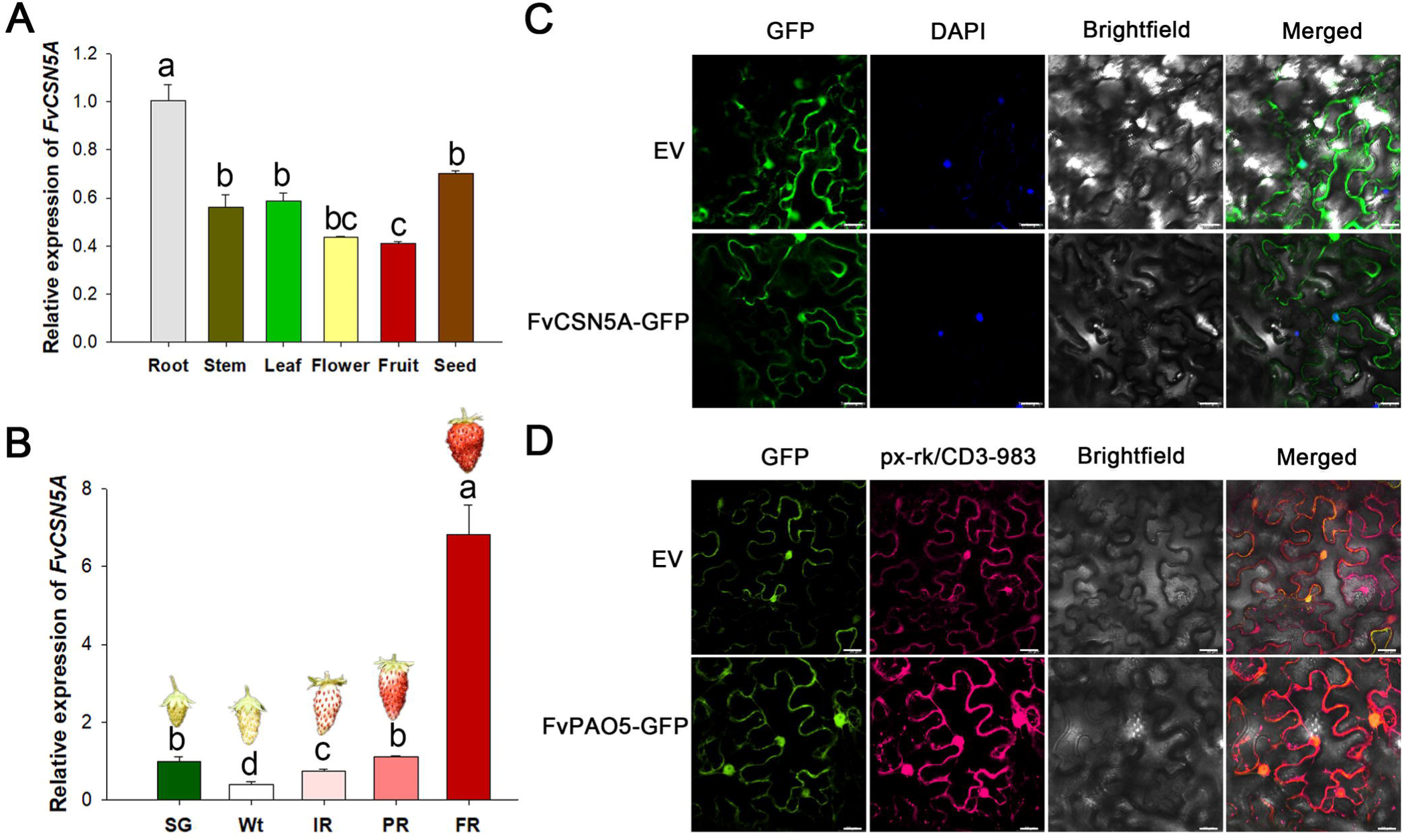
*FvCSN5A* expression pattern and subcellular localization. (**A)** *FvCSN5A* expression levels in different tissues of diploid strawberries (*F. vesca*, Ruegen). Different letters indicate statistically significant differences at P<0.05 as determined by one-way ANOVA. (**B)** *FvCSN5A* expression levels in fruit receptacles at different developmental stages of diploid strawberries (*F. vesca*, Ruegen). SG, small green fruit stage; Wt, white fruit stage; IR, initial red fruit stage; PR, partial red fruit stage; FR, full ripening fruit stage. Relative expression values were relative to receptacles at the SG stage, which was assigned an arbitrary value equal to one. Different letters indicate statistically significant differences at P<0.05 as determined by one-way ANOVA. (**C)** Subcellular localization of FvCSN5A-GFP fusions in transiently transformed *N. benthamiana* leaves. All experiments were performed 48 h post-infiltration. Bars, 20 μm. For the locations of the nuclei, *N. benthamiana* leaves were stained with DAPI. Px-rk/CD3-983 was a marker for the locations of the peroxisome.

To determine the subcellular localization of FvCSN5A, we expressed the Super1300:FvCSN5A-GFP construct or the empty vector (EV, expressing GFP alone) in tobacco leaves and performed fluorescence imaging observations. The results showed that the FvCSN5A-GFP signals were detectable in the cytoplasm and nucleus, the latter of which overlapped with the 4’,6-diamidino-2-phenylindole dihydrochloride (DAPI) staining, a dye used to label nuclei (Fig. 2C). In comparison, FvPAO5 was localized in the nucleus and peroxisomes, as evidenced by its co-localization with the peroxisome marker px-RK/CD3-983 (Fig. 2C). Overall, the common localization of the two proteins in the nucleus suggests their potential roles important to strawberry plant development.

### Inhibition of *FvCSN5A* expression led to developmental defects in leaves and flowers

CSN5 has been found to participate in crucial biological functions, such as development and stress responses in Arabidopsis and tomato (Shang et al., 2019; Qin et al., 2020; Lu et al., 2024); however, the function of strawberry homolog FvCSN5A remains unknown. To elucidate the role of FvCSN5A, we generated RNAi-based FvCSN5A-knockdown *F. vesca* plants. Two independent transgenic lines exhibiting different expression levels of *FvCSN5A* were obtained and verified by red fluorescence (Fig. S1). The expression levels of *FvCSN5A* in the RNAi-1 and RNAi-2 lines were decreased by approximately 8-fold and 46-fold, respectively (Fig. 3A). In addition, the chromosome ploidy of transgenic strawberry was analyzed by flow cytometry, and it was found that *FvCSN5A* RNAi-1 and RNAi-2 were both diploid strawberries (Fig. S2).

**Figure 3.**
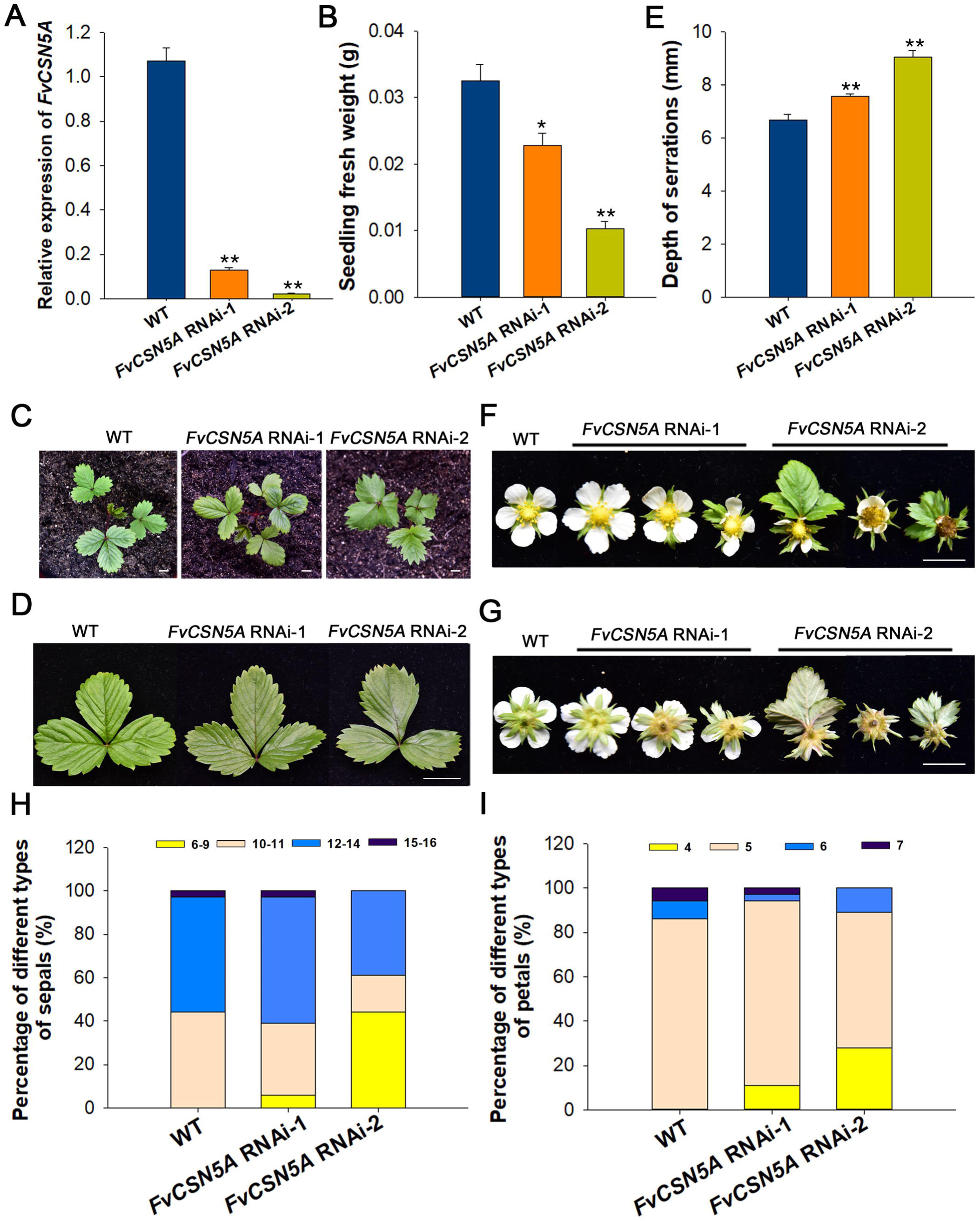
Phenotypes of the *FvCSN5A* RNAi transgenic plants. (**A)** Relative expression levels of *FvCSN5A* were determined by RT-qPCR. (**B)** The fresh weight of wild-type (WT, *F. vesca*, Ruegen) and *FvCSN5A* RNAi transgenic plants. The seedlings grown 45 days after germination were collected for the experiments. (**C)** The photos of WT and *FvCSN5A* RNAi transgenic plants in soil. Bars, 1 cm. (**D)** One compound leaf of WT and *FvCSN5A* RNAi transgenic plants. Bars, 1 cm. (**E)** Depth of the serrations of WT and *FvCSN5A* RNAi transgenic plants. (**F)** The flowers of WT and *FvCSN5A* RNAi transgenic plants. Bars, 1 cm. (**G)** The sepals of WT and *FvCSN5A* RNAi transgenic plants. Bars, 1 cm. (**H)** The percentages of different types of sepals in WT and *FvCSN5A* RNAi transgenic plants. n = 36. (**I)** The percentages of different types of petals in WT and *FvCSN5A* RNAi transgenic plants. n = 36. Statistical significance of one-way ANOVA: *, P < 0.05; **, P < 0.01.

The fresh weight of RNAi seedlings was smaller than that of the wild-type (Fig. 3B). The RNAi plants maintained a similar leaf shape index and number of serrations compared to the wild-type leaves (Fig. S3A-B). However, the leaves of *FvCSN5A* RNAi-1 and *FvCSN5A* RNAi-2 plants exhibited deeper serrations with sharper serrated blade angles (Fig. 3C-E). Moreover, the petal and sepal numbers of the RNAi plants exhibited significant variation. In the wild-type, each flower typically has 10 to 14 sepals and 5 petals (Fig. 3F-I). In contrast, some flowers of the *FvCSN5A* RNAi plants had 6 to 9 sepals (6% in RNAi-1 and 44% in RNAi-2) and 4 petals (11% in RNAi-1 and 28% in RNAi-2), while the wild-type plant did not show such variations (Fig. 3F-I). Collectively, the reduction in *FvCSN5A* expression resulted in substantial developmental defects in both leaves and flowers.

### FvCSN5A affected the fertility of strawberries, especially pollen viability

Since plant flowering and seed formation are fundamental to strawberry fruit development (Liao et al., 2018), we examined the influence of FvCSN5A on seed development. The *FvCSN5A* RNAi-1 and *FvCSN5A* RNAi-2 transgenic plants exhibited no observable defects in the appearance of the stigmas, carpels, or stamens (Fig. S3C-D). However, the number of carpels and anthers was reduced in *FvCSN5A* RNAi flowers (Fig. S3E-F).

Subsequently, we performed Alexander’s staining and in vitro pollen germination assays to assess pollen viability (Fig. 4A-B). Based on Alexander’s staining, we determined that there was near-complete pollen activity abortion in the lower expression line (*FvCSN5A* RNAi-2; 10.29±0.98) and a subtle reduction in the moderate expression line (*FvCSN5A* RNAi-1; 91.35±1.22; Fig. 4C). A significant decrease in pollen germination in vitro was observed in *FvCSN5A* RNAi-1 (65.93±7.07) and *FvCSN5A* RNAi-2 (8.49±2.42) compared to the wild-type (96.97±3.03; Fig. 4D). Furthermore, the germinated pollen tube length of *FvCSN5A* RNAi plants was shorter than that of the wild-type (Fig. 4E). These results demonstrate that the inhibition of *FvCSN5A* expression leads to reduced pollen viability in strawberry.

**Figure 4.**
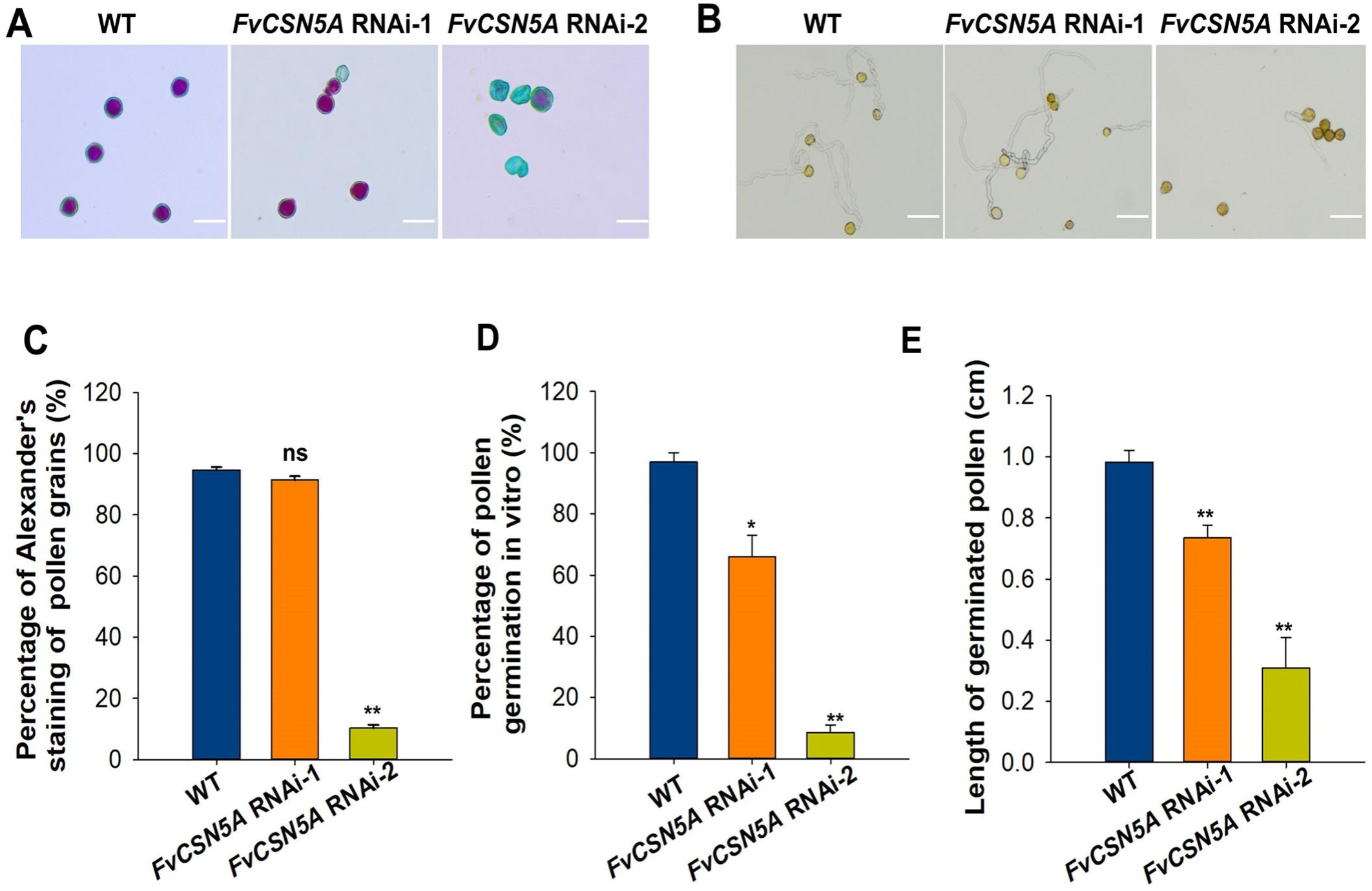
Pollen grain fertility in *FvCSN5A* RNAi transgenic plants. (**A)** The images of Alexander’s staining of pollen grains from WT and *FvCSN5A* RNAi transgenic plants. The red-stained pollen grains are viable and fertile. Bars, 20 μm. (**B)** The images of pollen tubes from WT and *FvCSN5A* RNAi transgenic plants germinating *in vitro*. Bars, 50 μm. (**C)** The percentage of Alexander’s staining of pollen grains from WT and *FvCSN5A* RNAi transgenic plants. (**D)** The percentage of pollen tubes from WT and *FvCSN5A* RNAi transgenic plants germinating *in vitro*. (**E)** The length of pollen tubes from WT and *FvCSN5A* RNAi transgenic plants germinating *in vitro*. Statistical significance of one-way ANOVA: *, P < 0.05; **, P < 0.01. Non-significant: ns

To further investigate the relative influence of male and female gametophytes on seed formation and fruit expansion, we performed reciprocal crosses between wild-type and *FvCSN5A* RNAi-2 plants. As shown in Fig. S4A, more seeds were produced, and fruit expanded to a greater extent when the wild-type pollen was applied to *FvCSN5A* RNAi-2 stigmas, compared to using *FvCSN5A* RNAi-2 as the paternal plant. Collectively, these data indicate that FvCSN5A is essential for strawberry reproductive success, with particular importance for pollen development.

### FvCSN5A positively regulated fruit ripening

To gain insight into the function of FvCSN5A on fruit ripening, we observed the fruit coloring phenotype of *FvCSN5A* RNAi. The results demonstrated that strawberry fruit coloration was retarded in *FvCSN5A* RNAi-1 plants (Fig. 5A), while *FvCSN5A* RNAi-2 fruits were impeded from completing the normal development and maturation process (Fig. S4B). The content of anthocyanin was lower in *FvCSN5A* RNAi-1 fruits compared to the control, consistent with the observed phenotype (Fig. 5B). The firmness analysis also revealed that the softening process was inhibited in *FvCSN5A* RNAi fruits (Fig. 5C). The fresh weight was further determined, indicating that the fresh weight of *FvCSN5A* RNAi fruits was smaller than that of the control (Fig. 5D). The contents of sucrose, glucose, fructose, and total soluble sugar were significantly lower in the fruit of *FvCSN5A* RNAi (Fig. 5E).

**Figure 5.**
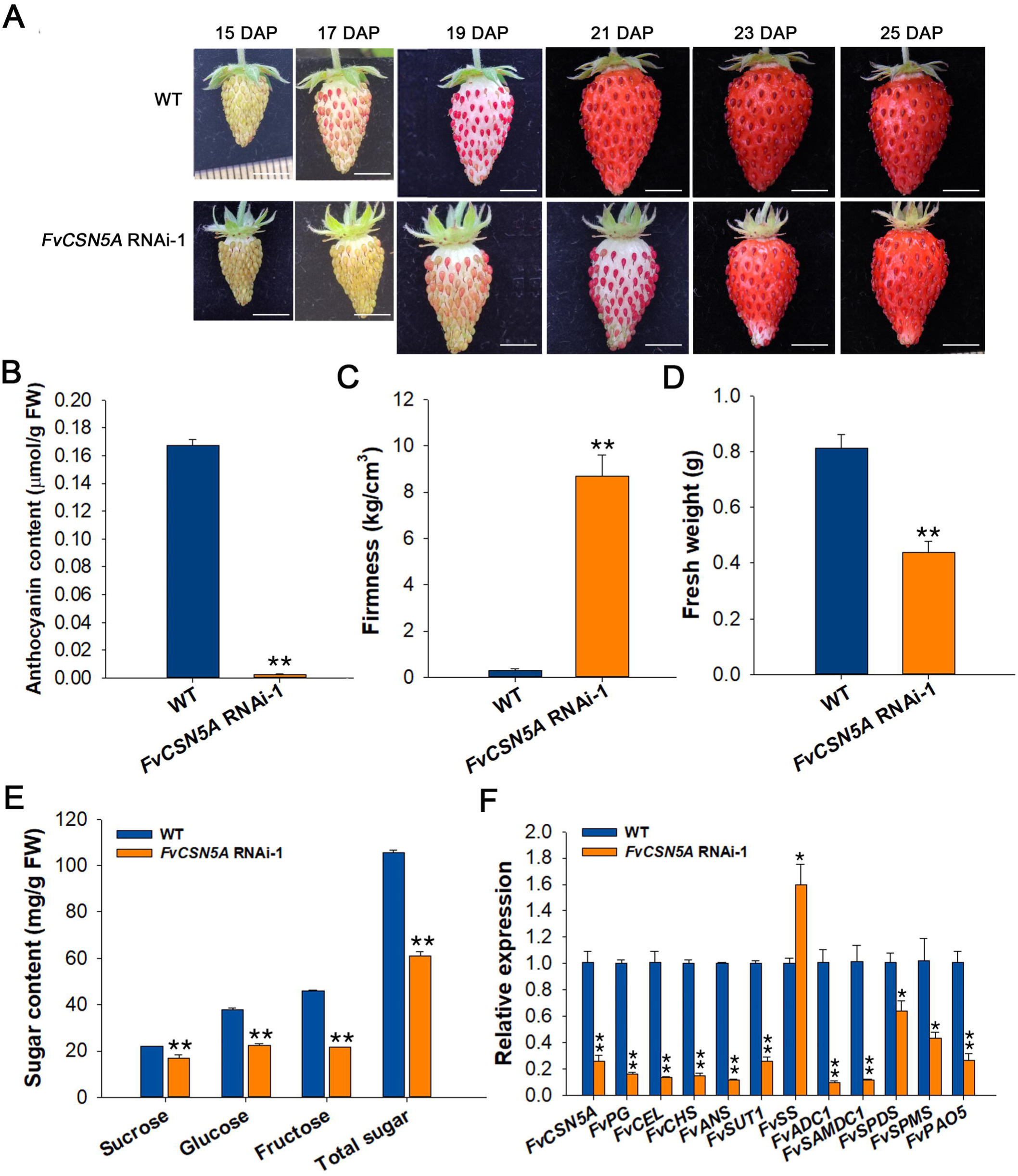
Fruit ripening changes in *FvCSN5A* transgenic fruits. (**A)** Photos were taken at 15, 17, 19, 21, 23, and 25 days after pollination (DAP) respectively to record the fruit phenotype. Bars, 0.5 cm. (**B-E)** The anthocyanin content **(B)**, firmness **(C)**, fresh weight **(D)**, and soluble sugar content **(E)** of WT and *FvCSN5A* RNAi transgenic fruits. Fruits were collected for experiments 21 days after pollination (DAP). Bars are means SEs of three independent experiments. (**F)** RT-qPCR was used to analyze the genes’ expression in WT and *FvCSN5A* RNAi transgenic fruits, and three biological replicates were tested for all samples using the *Actin* gene from strawberries as an internal control. The relative expression levels were calculated using the 2^-△△Ct^ method. FvCSN5A: COP9 Signalosome5; FvPG: polygalacturonase; FvCEL: cellulose; FvCHS: chalcone synthase; FvANS: anthocyanidin synthase; FvSUT1: sucrose transporter1; FvSS: sucrose synthase. FvADC1: arginine decarboxylase1; FvSAMDC1: S-adenosyl-methionine decarboxylase1; FvSPDS: spermidine synthase; FvSPMS: spermine synthase; FvPAO5: polyamine oxidase5. Statistical significance was determined by one-way ANOVA: *, P < 0.05; **, P < 0.01.

The expression levels of genes related to sugar and anthocyanin accumulation, fruit softening, and PA homeostasis in transgenic strawberry fruits were also examined using RT-qPCR. The expression levels of sucrose transporter *FvSUT1*, chalcone synthase *FvCHS* and anthocyanidin synthase *FvANS*, polygalacturonase *FvPG* and cellulose *FvCEL* were significantly inhibited in the RNAi fruit, while sucrose synthase *FvSS*, which negatively regulates sugar accumulation, was elevated (Fig. 5F). Based on the expression of these marker genes, the physiological results, and phenotypic observations, these findings indicate that FvCSN5A regulated fruit ripening and fruit quality.

To explore the effect of FvCSN5A on PA metabolism, the relative gene expression levels in the WT and *FvCSN5A* RNAi fruits were detected. The results showed that the expression levels of polyamine synthetase (*FvADC1*, *FvSAMDC1*, *FvSPDS*, and *FvSPMS*) and polyamine oxidase (*FvPAO5*) in *FvCSN5A* RNAi fruits were significantly lower than that in WT. At the same time, we found that in the *FvCSN5A* RNAi-1 fruit, the expression level of all COP9 signalosome subunits, except *FvCSN1*, was significantly reduced (Fig. S3G).

To further verify the role of strawberry CSN5A, we manipulated *FaCSN5A* expression in octoploid cultivated strawberry fruits by overexpression or RNAi construction through transient transformation. The phenotype and RT-qPCR analysis demonstrated that the fruit with lower *FaCSN5A* mRNA levels (*FaCSN5A* RNAi) colored slowly compared to the control fruit, while the fruit with higher *FaCSN5A* mRNA levels (*FaCSN5A* OE) colored rapidly (Fig. S5A-B). The anthocyanin, soluble sugar, firmness and their relative gene expression were detected and significantly changed in *FaCSN5A* OE and RNAi fruits (Fig. S5C-D). We also examined the expression of PA metabolism-related genes. The results showed that the expression levels of *FaADC1*, *FaSAMDC1*, *FaSPDS*, *FaSPMS*, and *FaPAO5* in the *FaCSN5A* RNAi fruit were significantly lower than those in the WT. The expression levels of these genes in the *FaCSN5A* OE fruit were significantly higher than those in the WT (Fig. S5D). Altogether, these results demonstrate that strawberry CSN5A controls fruit ripening as a positive regulator at both physiological and molecular levels.

### FvCSN5A promoted FvPAO5 ubiquitination degradation

The CSN (COP9 signalosome) complex typically regulates plant development and stress response by modulating the E3 ubiquitin ligase complex. Therefore, we propose that the FvCSN5A-FvPAO5 interaction might affect FvPAO5 protein stability and regulate strawberry development and fruit ripening.

Firstly, we examined the protein levels of FvPAO5 and FvCSN5A at different stages of fruit development. The results showed a decrease in the levels of FvPAO5 during fruit ripening, while the levels of FvCSN5A increased gradually (Fig. 6A). Additionally, FvPAO5 exhibited relatively higher accumulation in the *FvCSN5A* RNAi plants compared to the WT plant (Fig. 6B). Secondly, we investigated the degradation pathway of FvPAO5 through the 26S proteasome in a cell-free system. The results showed that the protein level of FvPAO5 decreased when incubated with strawberry total protein, and this degradation was inhibited by the proteasome inhibitor MG132 (Fig. 6C). These data indicate that FvCSN5A may promote the degradation of FvPAO5.

**Figure 6.**
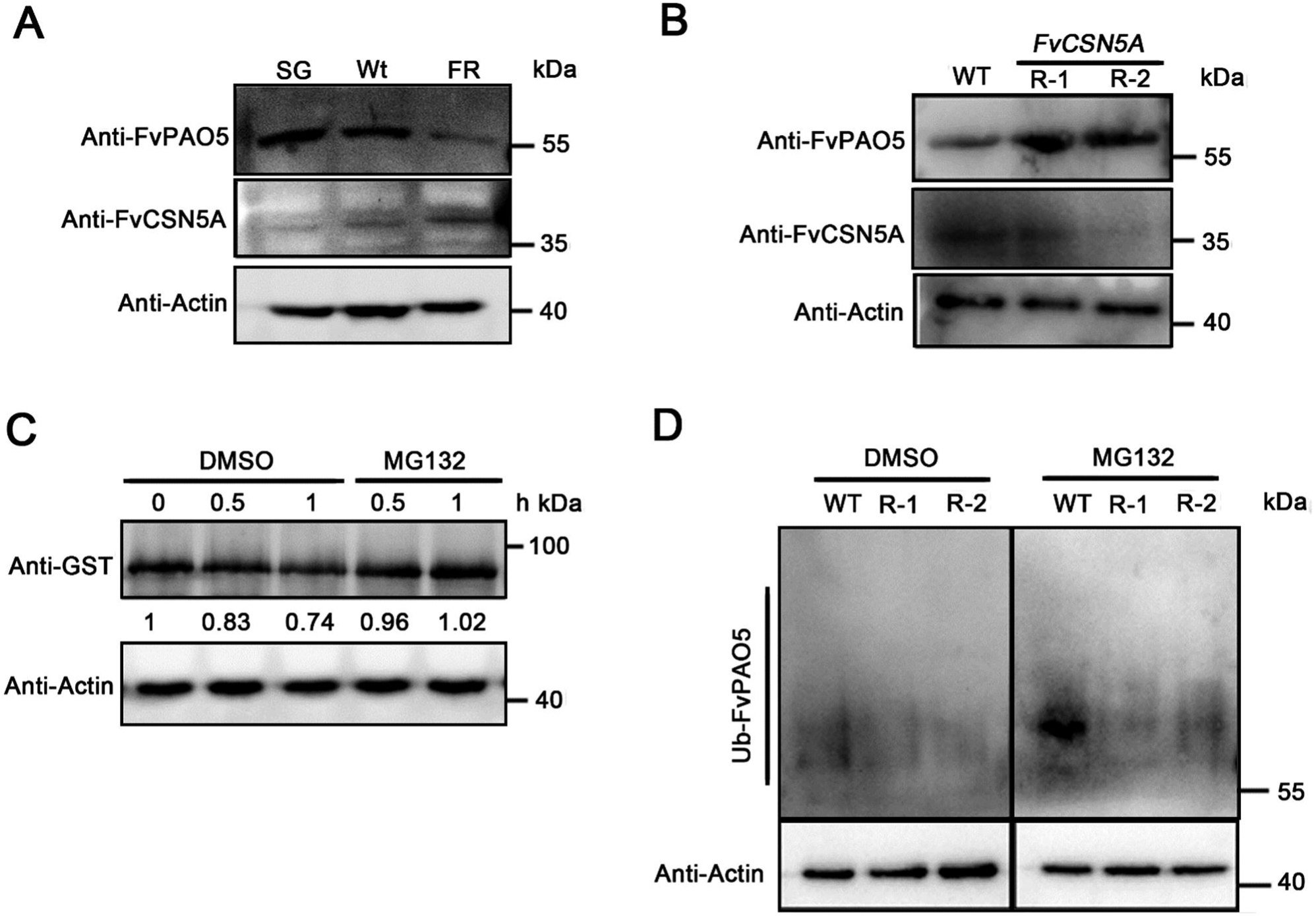
The ubiquitination degradation of FvPAO5 is dependent on FvCSN5A. (**A)** FvPAO5 and FvCSN5A protein levels in fruit receptacles at different developmental stages of diploid strawberry (*F. vesca*, Ruegen). SG, small green fruit stage; Wt, white fruit stage; FR, full ripening fruit stage. (**B)** FvPAO5 and FvCSN5A protein abundance detection in the leaves from *FvCSN5A* RNAi and WT plants. FvPAO5 and FvCSN5A were detected by anti-FvPAO5 and anti-FvCSN5A, respectively. (**C)** Cell-free FvPAO5 protein degradation. Total protein was extracted from strawberry leaves and incubated with FvPAO5-GST protein for 0.5, and 1 h with MG132 treatment or DMSO treatment as a control. FvPAO5-GST protein and total plant protein were detected by anti-GST and anti-Actin, respectively. (**D)** *In vivo* FvPAO5 polyubiquitination in *FvCSN5A* RNAi and WT plants. Three-month-old then harvested for protein extraction. Each total protein was incubated with P62-agarose to obtain ubiquitinated proteins. Polyubiquitinated FvPAO5 was detected with an anti-FvPAO5 antibody. The accumulation of Actin in the total protein was tested as a control.

The polyubiquitination of FvPAO5 was assessed in the *FvCSN5A* RNAi and WT plants. Tissue culture seedlings of the *FvCSN5A* RNAi and WT plants were treated with or without 50 μM MG132 for 5 h and then harvested for protein extraction. The ubiquitination-modified proteins were enriched with P62-agarose, and ubiquitinated FvPAO5 was detected by an anti-FvPAO5 antibody. Without MG132 treatment, polyubiquitinated FvPAO5 was weak in the WT plants, while it appeared significantly stronger under MG132 treatment. However, the FvPAO5 polyubiquitination signal was hardly visible in the *FvCSN5A* RNAi plants, regardless of whether MG132 was used or not (Fig. 6D).

These data suggest that FvCSN5A may promote the proteasome-dependent degradation of FvPAO5 in a ubiquitin-mediated manner.

### FvCSN5A interacted with FvCUL1 in vitro and in vivo

Many studies have reported that CSN interacts with CUL within the E3 ubiquitin ligase complex, and the deneddylation of AtCUL1 was blocked in the *atcsn5* mutant (Schwechheimer et al., 2001; Cope and Deshaies, 2003; Gusmaroli et al., 2004). To determine whether FvCSN5A, FvCULs, and FvPAO5 function within the same complex, we performed a CoIP mass spectrometry assay with an anti-FvPAO5 antibody. The results showed that, in addition to FvCSN5A, FvCSN1 and FvCUL1 were also detected in the anti-FvPAO5 antibody-enriched proteins from the WT (Supplemental Data Set 1). However, FvPAO5 and FvCUL1 did not interact directly in the yeast two-hybrid experiment (data not shown). This implies that FvPAO5 might associate with FvCUL1 via FvCSN5A. To examine the relationship between FvCSN5A and FvCUL1, we investigated the interaction between FvCUL1 and FvCSN5A. The results showed that FvCUL1 interacts with FvCSN5A in Y2H and FLC assays (Fig. 7A-B). Therefore, we hypothesize that FvCSN5A may collaborate with FvCUL1 to regulate FvPAO5 protein stability.

**Figure 7.**
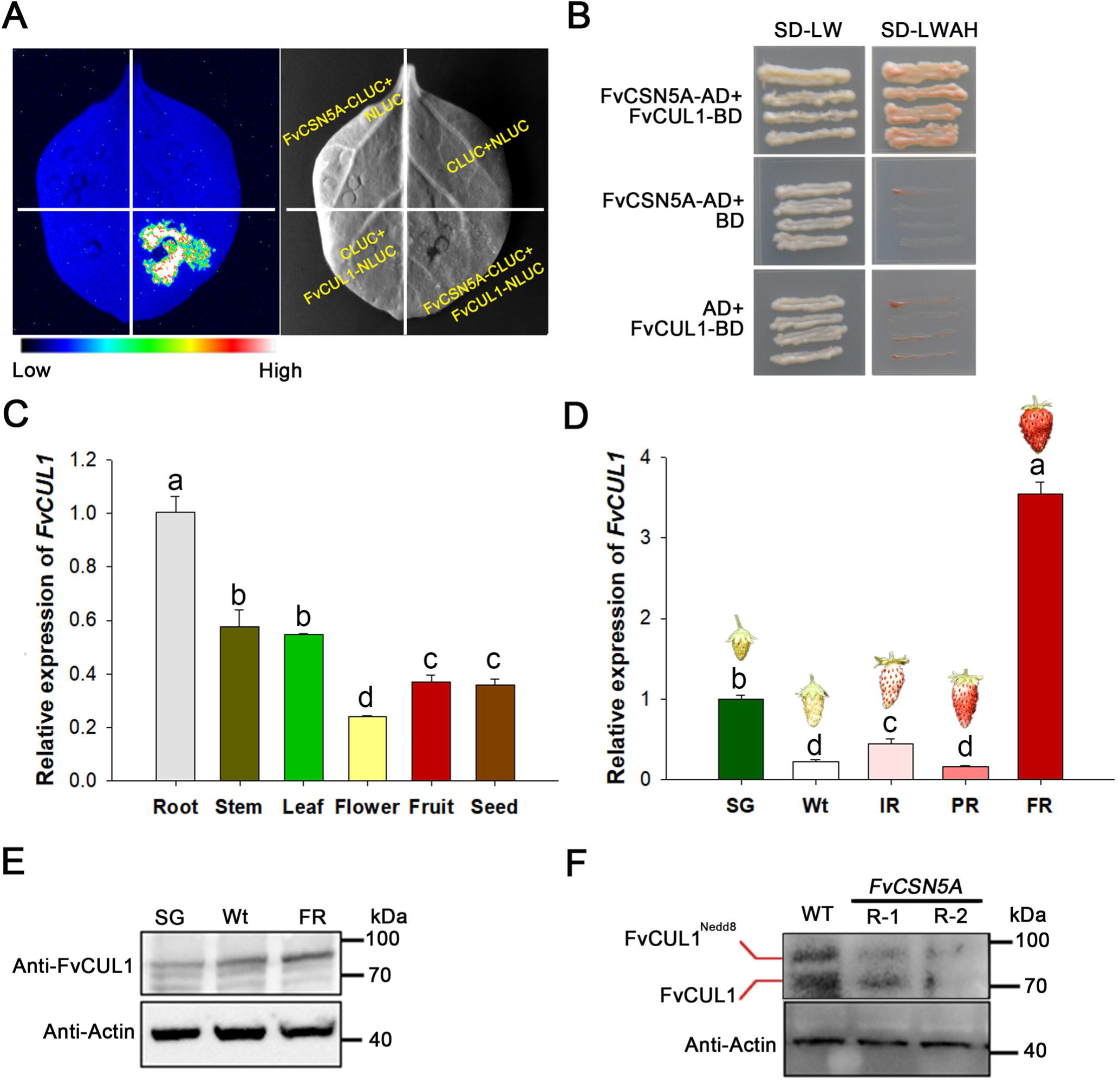
FvCUL1 interacted with FvCSN5A *in vitro* and *in vivo*. (**A)** FvCUL1 interacted with FvCSN5A in FLC assay. The CDS of *FvCSN5A* was constructed into pCAMBIA1300-CLUC vector, and the CDS of *FvCUL1* was constructed into pCAMBIA1300-NLUC vector. (**B)** FvCUL1 interacted with FvCSN5A in yeast two-hybrid assay. SD-LW: synthetic dropout medium without Leu and Trp; SD-LWHA: synthetic dropout medium without Leu, Trp, His, and Ade. (**C)** The transcript level of *FvCUL1* in different tissues of strawberry. Different letters indicate statistically significant differences at P<0.05 as determined by one-way ANOVA. (**D)** *FvCUL1* transcript levels in fruit receptacles at different developmental stages of diploid strawberry. Different letters indicate statistically significant differences at P<0.05 as determined by one-way ANOVA. (**E)** FvCUL1 protein level in fruit receptacles at different developmental stages of diploid strawberries. (**F)** Detection of FvCUL1 protein level in *FvCSN5A* RNAi and WT plants using antibody anti-CUL1. FvCUL1^Nedd8^: FvCUL1 with neddylation; FvCUL1: FvCUL1 without neddylation.

In strawberries, the *FvCUL1* transcript level was highest in the root, and lowest in flowers (Fig. 7C). During fruit ripening, the expression pattern of *FvCUL1* was similar to *FvCSN5A*, rising rapidly and reaching its peak at the FR stage in both mRNA and protein levels (Fig. 7D-E). Subcellular localization analysis was performed and found that FvCUL1 was localized in the cytoplasm (Fig. S6). We also tested the influence of FvCSN5A on the FvCUL1 protein level using an anti-CUL1 antibody. We found the deneddylated and neddylated forms of FvCUL1 in the wild-type, and the FvCUL1 protein level in the *FvCSN5A* RNAi line markedly decreased (Fig. 7F). These results suggested that FvCSN5A, the subunit of CSN, interacts with FvCUL1, which may affect the protein stability of FvCUL1.

### Polyamine, phytohormone, H_2_O_2_ and Dap levels in the *FvCSN5A* RNAi plants

Previous studies have reported that CSN5A is involved in numerous phytohormones, including indoleacetic acid (IAA), jasmonic acid (JA), and gibberellic acid (GA; Jin et al., 2014; Qin et al., 2020). In the current investigation, we have determined that FvCSN5A promotes the ubiquitin-mediated degradation of FvPAO5, a key enzyme that catalyzes the conversion of Spd/Spm to H_2_O_2_ and 1,3-diaminopropane (Dap; Mo et al., 2020). To elucidate the regulatory roles of FvCSN5A, we assessed the alterations in endogenous PAs, phytohormones, H_2_O_2_, and Dap in the *FvCSN5A* RNAi plant.

Firstly, we quantified PA levels in strawberries. The contents of Spd and Spm were significantly lower in the *FvCSN5A* RNAi plant compared to the WT plant, with Spd being almost undetectable in the *FvCSN5A* RNAi-2 plant (Fig. 8A-B). However, Put accumulation varied, showing a reduction in in the *FvCSN5A* RNAi-1 line but not significantly different from WT in the *FvCSN5A* RNAi-2 line (Fig. 8C). These results indicate that the inhibition of strawberry *FvCSN5A* expression leads to a substantial decrease in Spd and Spm levels, which may be attributable to the over-accumulation of the FvPAO5 protein.

**Figure 8.**
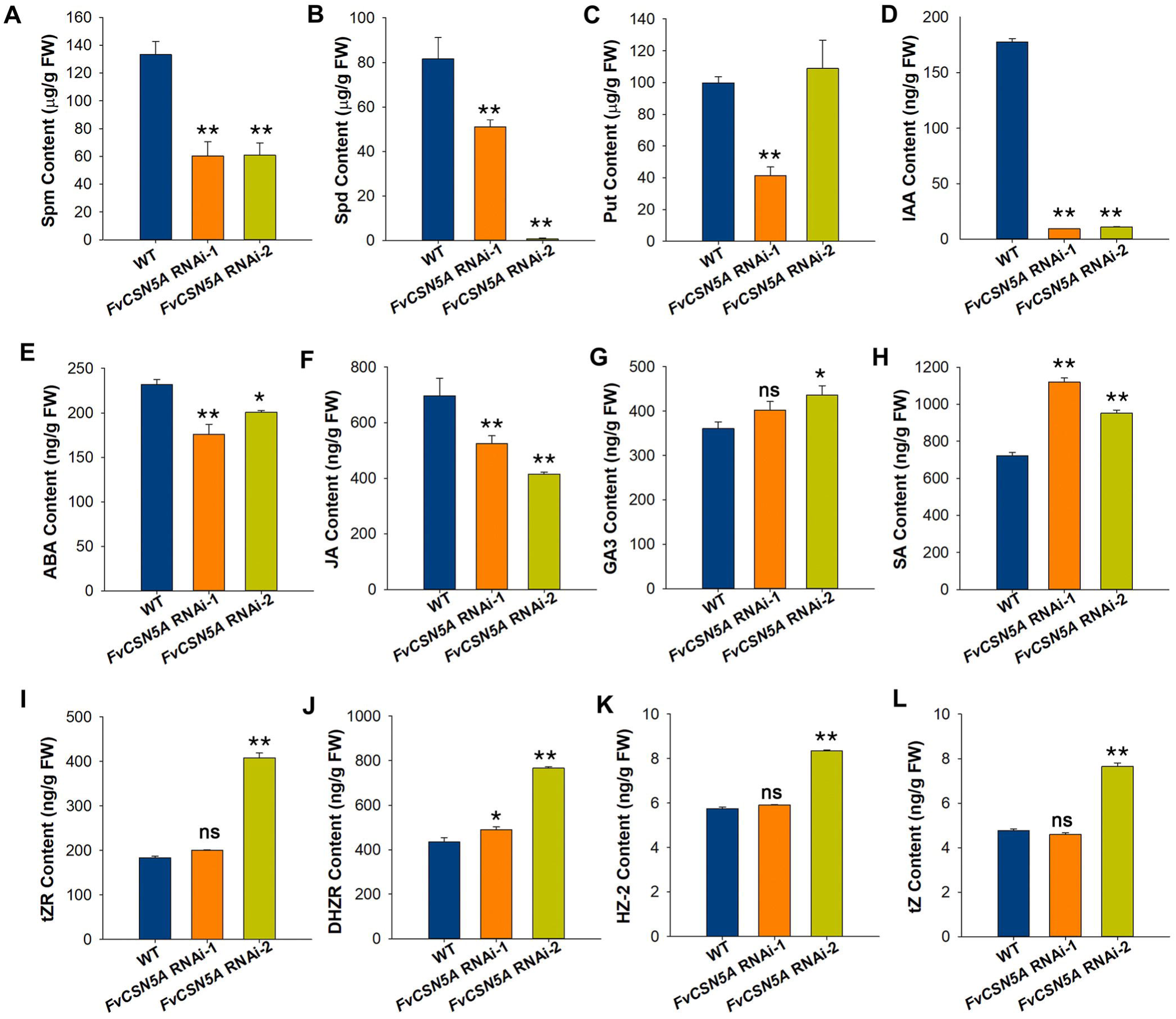
Polyamine and phytohormone contents in *FvCSN5A* RNAi transgenic plants (**A-C)** Spm **(A)**, Spd **(B)**, and Put **(C)** content in WT and *FvCSN5A* RNAi transgenic leaves. (**D-L)** Phytohormone IAA, ABA, JA, GA3, SA, tZR, DHZR, HZ-2 and tZ content in WT and *FvCSN5A* RNAi transgenic leaves. Statistical significance of one-way ANOVA: *, P < 0.05; **, P < 0.01. Non-significant: ns

Secondly, we investigated the contents of phytohormones related to plant development and environmental responses, including IAA, ABA, JA, GA, cytokinin (CK), and salicylic acid (SA). The synthesis of IAA, ABA, and JA was suppressed in *FvCSN5A* RNAi plants, with IAA displaying an 83% reduction (Fig. 8D-F). In strawberries, the most abundant GA compound was GA3, and the predominant CK compounds were TZR and DHZR, especially, the levels of GA3, TZR, DHZR, and SA were elevated in *FvCSN5A* RNAi plants (Fig. 8G-I).

At last, we assessed the amount of H_2_O_2_ and Dap in the *FvCSN5A* RNAi plants. The DAB staining results showed that the pollen grains, leaves and roots of *FvCSN5A* RNAi plants, particularly *FvCSN5A* RNAi-2, were stained more intensely, indicating higher H_2_O_2_ levels compared to the lighter staining observed in the WT (Fig. 9A-B, Fig. S7). Furthermore, the ROS generation, as detected by 2,7□dichlorodihydrofluorescein diacetate (DCFH□DA) fluorescence, was markedly higher in the pollen grains and leaves of *FvCSN5A* RNAi plants (Fig. 9C-D). The content analysis also confirmed that the *FvCSN5A* RNAi plant had significantly elevated H_2_O_2_ production compared to the WT (Fig. 9E). The content of Dap, another product of the FvPAO5-catalyzed reaction, was significantly higher in the *FvCSN5A* RNAi plant compared to the WT (Fig. 9F). Moreover, when strawberries were treated with exogenous Spd, the Dap content in *FvCSN5A* RNAi plants decreased, while it increased in the WT (Fig. 9F), reducing the gap between the two.

**Figure 9.**
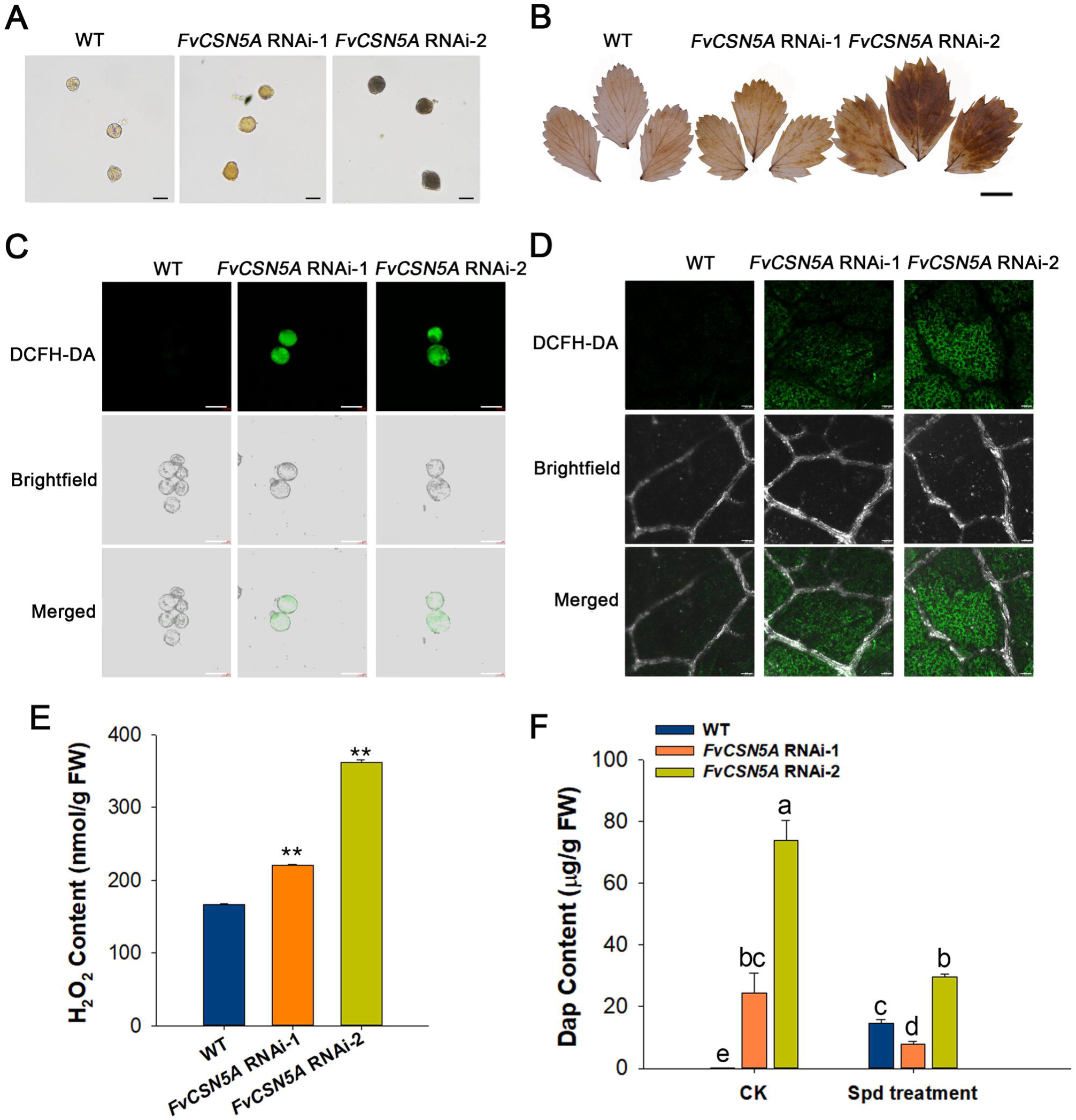
Detection of ROS and Dap accumulation in *FvCSN5A* RNAi transgenic plants (**A)** DAB dye of pollen grains from WT and *FvCSN5A* RNAi transgenic plants. Bars, 20 μm. (**C)** ROS signal determination of pollen grains by DCFH-DA. Bars, 20 μm. (**D)** ROS signal determination of leaves by DCFH-DA. Bars, 50 μm. (**E)** H_2_O_2_ content in WT and *FvCSN5A* RNAi transgenic plants. (**F)** Dap content in WT and *FvCSN5A* RNAi transgenic plants after Spd treatment. Statistical significance of one-way ANOVA: *, P < 0.05; **, P < 0.01.

Overall, these results indicate that the reduction of *FvCSN5A* expression leads to a significant decrease in the contents of PAs and IAA, accompanied by an excessive accumulation of H_2_O_2_ and Dap. The disruption of this homeostatic balance in *FvCSN5A* RNAi plants may contribute to the

### Overexpression of *FvPAO5* affected plant development and pollen viability

We previously found that FaPAO5 negatively regulated fruit ripening (Mo et al., 2020). The above results suggest that FvCSN5A plays a role in plant development and fruit ripening by affecting FvPAO5 stability. To determine which functions of FvCSN5A are mediated by FvPAO5, we produced and characterized *FvPAO5* OE lines using Super1300:GFP (empty vector, EV) and Super1300:FvPAO5-GFP (*FvPAO5* OE) transgenic strawberry plants via the leaf disk transformation method with GFP as a selection tag (Fig. S8A-C). Two overexpression transgenic lines with different expression levels were used for subsequent experiments (Fig. 10A). Subcellular localization experiments suggested that FvPAO5 is expressed in both the cytoplasm and nucleus of *FvPAO5* OE plants, which was consistent with previous results (Fig. 10B).

**Figure 10.**
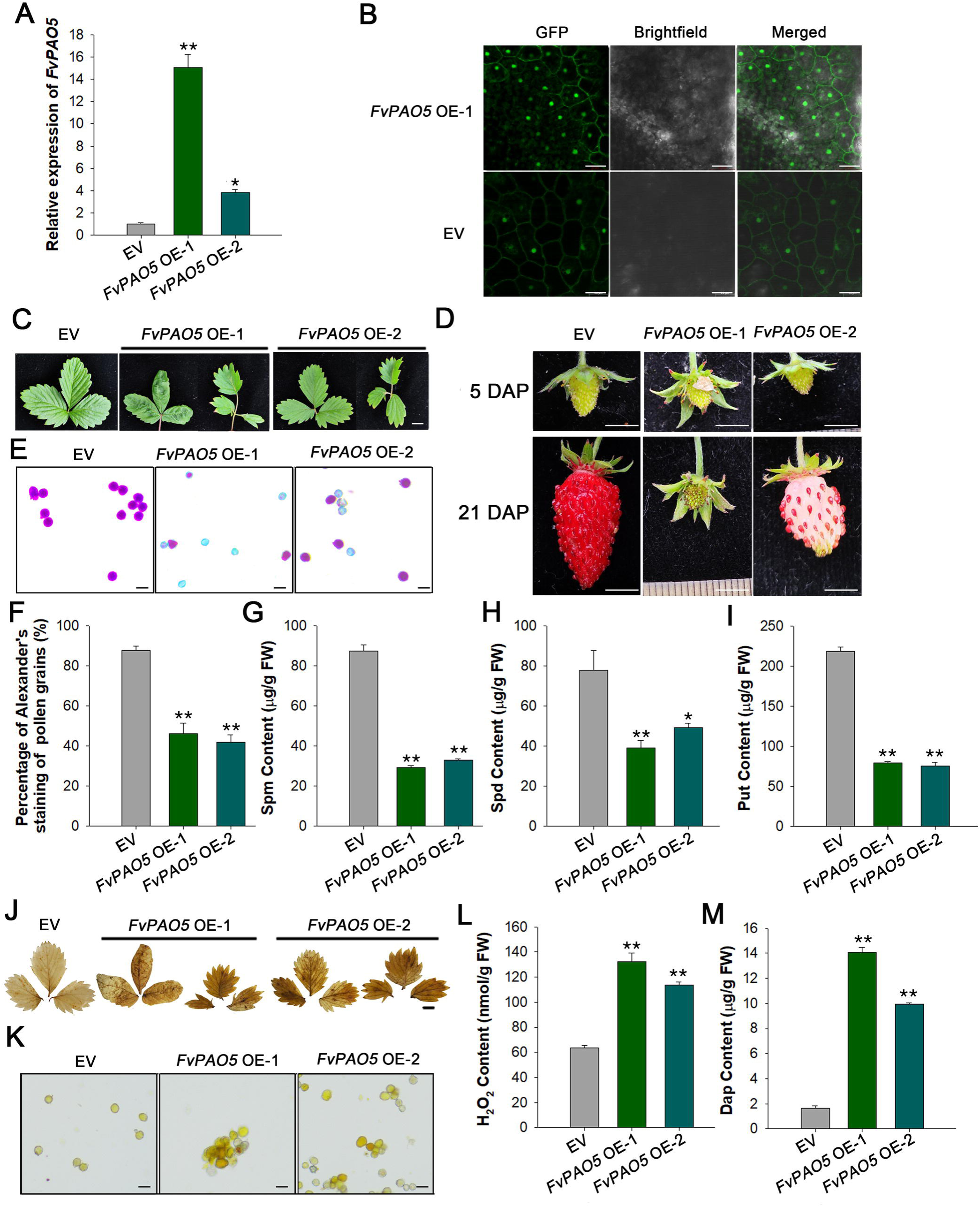
Phenotypes of the *FvPAO5* overexpression transgenic plants. (**A)** Relative expression levels of *FvPAO5* were determined in *FvPAO5* overexpression (OE) transgenic plants by RT-qPCR. EV: Super1300:GFP; *FvPAO5* OE: Super1300:FvPAO5-GFP. (**B)** Subcellular localization of FvPAO5 in *FvPAO5* OE transgenic plant leaves mesophyll cells. Bars, 20 μm. (**C)** Compound leaves in WT and *FvPAO5* OE transgenic plants. Bars, 1 cm. (**D)** Fruit development and ripening process in WT and *FvPAO5* OE transgenic plants. Bars, 0.5 cm. Photos were taken at 5 and 21 days after pollination respectively to record the fruit phenotype. (**E)** The images showing Alexander’s staining of pollen grains in WT and *FvPAO5* OE transgenic plants. The red-stained pollen grains are viable and fertile. (**F)** The percentages of Alexander’s staining of pollen grains in WT and *FvPAO5* OE transgenic plants. (**G-I)**: Spm **(G)**, Spd **(H)**, and Put **(I)** content in WT and *FvPAO5* OE transgenic leaves. (**J)** DAB dye of pollen grains in WT and *FvPAO5* OE transgenic plants. Bars, 20 μm. (**K)** DAB dye of one compound leaf in WT and *FvPAO5* OE transgenic plants. Bars, 1 cm. (**L)** H_2_O_2_ content in WT and *FvPAO5* OE transgenic plants. (**M)** Dap content in WT and *FvPAO5* OE transgenic plants. Statistical significance of one-way ANOVA: *, P < 0.05; **, P < 0.01.

The leaves of *FvPAO5* OE-1 and *FvPAO5* OE-2 plants exhibited different leaf shapes, numbers, and depths of serrations (Fig. 10C), with no significant differences in flower organ development between *FvPAO5* OE and EV plants (Fig. S7C). Observations of the fruit developmental process in *FvPAO5* OE plants revealed that the fruits of *FvPAO5* OE were deformed and fruit ripening was inhibited (Fig. 10D). Alexander’s staining showed a significant reduction in pollen activity in the *FvPAO5* OE lines (Fig. 10E-F). PAs and products of the FvPAO5-catalyzed reaction, including H_2_O_2_and Dap, were determined in the *FvPAO5* OE plants. The results showed significantly lower PA contents in the *FvPAO5* OE plant compared to the EV plant (Fig. 10G-I), while the accumulation of H_2_O_2_ and Dap was greater in *FvPAO5* OE plants than in EV plants (Fig. 10J-M). These experiments indicate that the overexpression of *FvPAO5* also affect plant development and pollen viability, similar to the phenotype observed in *FvCSN5A* RNAi plants, suggesting a functional association between FvCSN5A and FvPAO5.

### FaCSN5A regulates fruit ripening and H_2_O_2_ homeostasis partly depending on FaPAO5

To explore the genetic interactions between strawberry CSN5A and PAO5, we conducted a transient transgenic fruit experiment to obtain *FaCSN5A* and *FaPAO5* co-RNAi strawberry fruits (*FaCSN5A* RNAi/*FaPAO5* RNAi). Agrobacterium containing *FaCSN5A* RNAi or *FaPAO5* RNAi vectors were injected into strawberry fruit individually or in combination, and photos were taken at 0, 3, 6, and 7 days after injection. The results showed that fruit coloring was inhibited in the *FaCSN5A* RNAi fruit, while it was promoted in the *FaPAO5* RNAi fruit (Fig. 11A-B). The *FaCSN5A* RNAi/*FaPAO5* RNAi fruits colored more rapidly than the *FaCSN5A* RNAi fruits (Fig. 11A-B). The content of anthocyanin was similar to the observed phenotype (Fig. 11C). Additionally, the H_2_O_2_ accumulation was also measured, revealing that the content of H_2_O_2_ in *FaCSN5A* RNAi/*FaPAO5* RNAi fruits was lower than *FaCSN5A* RNAi fruits but higher than that in *FaPAO5* RNAi fruits (Fig. 11D). These findings suggest that FaCSN5A regulates fruit ripening and H_2_O_2_ homeostasis, partly depending on FaPAO5.

**Figure 11.**
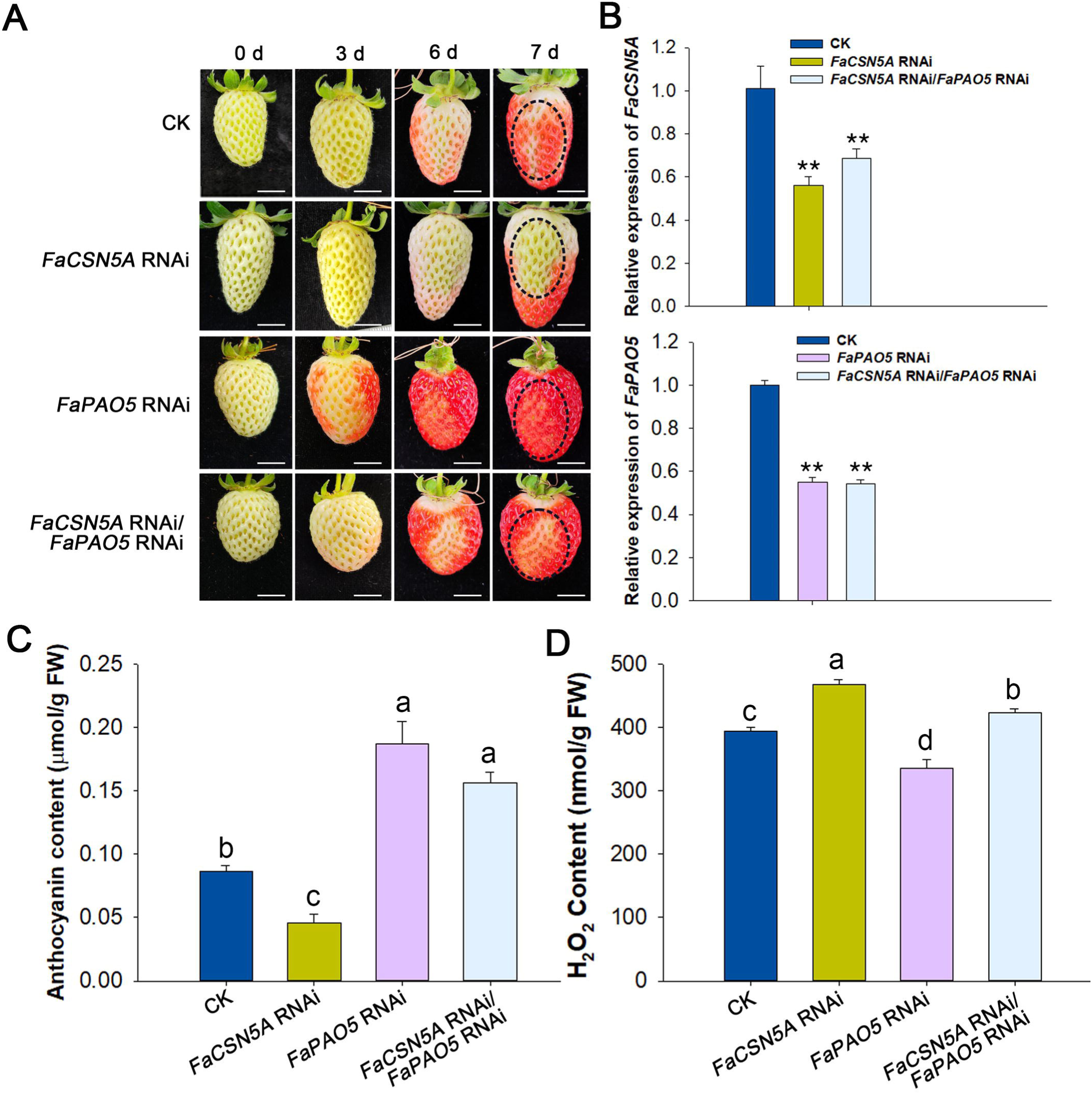
FaCSN5A control fruit ripening and H_2_O_2_ homeostasis partly depending on FaPAO5. (**A)** Phenotypes of *FaCSN5A* RNAi, *FaPAO5* RNAi, and *FaCSN5A* RNAi/*FaPAO5* RNAi strawberry fruits. Agrobacterium GV3101 strains containing recombinant plasmids were injected into DG fruits attached to the plant. The fruit phenotype was recorded at 0, 3, 6, and 7 days after injection. Bars, 0.5 cm. (**B)** Analysis of *FaCSN5A* and *FaPAO5* transcript levels in *FaCSN5A* RNAi, *FaPAO5* RNAi, and *FaCSN5A* RNAi/*FaPAO5* RNAi strawberry fruits. Fruits were collected for experiments 7 days after injection. (**C-D)** The anthocyanin content **(C)** and H_2_O_2_ content **(D)** of *FaCSN5A* RNAi, *FaPAO5* RNAi, and *FaCSN5A* RNAi/*FaPAO5* RNAi strawberry fruits. Fruits were collected for experiments 7 days after injection. Statistical significance of one-way ANOVA: *, P < 0.05; **, P < 0.01; different letters indicate statistically significant differences at P<0.05.

## DISCUSSION

Strawberries, as a perennial herbaceous plant, has emerged as a model organism for investigating the non-climacteric fruit ripening process, regulated by the intricate crosstalk of PAs with ABA, IAA, and ethylene (Guo et al., 2018; Bai et al., 2021; Gao et al., 2021). Consistent with the earlier notion of "no growth, no PA production" (Russell, 1973), the present data not only further corroborate the pivotal role of PAs in fruit ripening, but also provide novel insights into the efficacy of PAs in modulating plant growth, flowering, as well as fruit yield and quality in strawberries, especially by the CSN5-PAO5 module in strawberries. This underscores the significance of the COP9 signalosome-ubiquitination system in PA metabolism, underlying plant vegetative growth, reproductive growth, and ultimately fruit ripening and quality.

### Spd/Spm plays essential roles in strawberry vegetative and reproductive growth

Our previous investigations have demonstrated that during strawberry fruit development, Put content gradually declined, reaching its nadir in the ripe fruit, while spermidine (Spd) content remained at low levels, albeit exhibiting a slight rise during fruit ripening (Guo et al., 2018). Remarkably, spermine (Spm) content rapidly increased at the onset of ripening, becoming a dominant component of PAs in the ripe fruit. This proves an important role of Spd/Spm, particularly Spm, confirmed by the functional analysis of FaSAMDC1, a key enzyme for Spd/Spm biosynthesis (Guo et al., 2018). Moreover, the protein expression levels of FaPAO5 gradually decreased during fruit development, indicating its function as a negative regulator of ripening, in line with observed the function of FaPAO5 and H_2_O_2_ levels (Mo et al., 2020). These studies uncover that elevated FaSAMDC1 and depressed FaPAO5 levels during fruit ripening synergistically promote Spd/Spm accumulation and regulate H_2_O_2_ homeostasis, suggesting a cooperative interplay between PAs and ROS. Indeed, it was found that the expression of *OsPAO5* in rice seedlings is induced by light and suppressed by darkness, with *ospao5* mutants exhibiting lower H_2_O_2_ levels, resulting in a longer mesocotyl, faster seedling growth, and improved crop yield (Lv et al., 2021). However, the post-translational regulation of PAO5 remains elusive.

Fortunately, an interactor of FvPAO5, FvCNS5A, has now been identified through comprehensive physiological, biochemical, and molecular evidence. Notably, *FvCNS5A* RNAi plants exhibited elevated levels of both FvPAO5 and H_2_O_2_ (Fig. 6 and 9) and reduced amounts of IAA, ABA, and JA (Fig. 8), ultimately impairing plant growth and flowering, as well as fruit yield and ripening (Fig. 3-5). Intriguingly, the contents of Spd and Spm were significantly lower in the RNAi plants (Fig. 8), while Put levels remained unchanged (Fig. 8). Given that strawberry PAO5 acts as a negative regulator of ripening by catalyzing the degradation of Spm/Spd in the TC-type (Mo et al., 2020), thus the stable Put accumulation is not derived from Spd/Spm degradation. The downregulation of both *ADC1* and *SAMDC1* expression in the RNAi plants (Fig. S5) suggests that the stable Put level might result from the inhibited Spd biosynthesis, while the PAO5-mediated degradation may function as a feedback control mechanism, potentially via H_2_O_2_. To a large extent, the homeostasis and balance among Put, Spd, and Spm as well as H_2_O_2_ are essential for strawberry vegetative and reproductive growth. When this balance is disrupted by abundant FvPAO5 in the *FvCSN5A* RNAi plants, the conversion of Put to Spm is blocked as a sign of the undetectable Spd, ultimately leading to abnormal plant development, particularly in the reproductive phase. Based on our current data and a recent report that the regulation of PA composition, especially the Put/Spm ratio, may be an effective strategy to enhance plant survival and fitness (Navakoudis and Kotzabasis, 2022), we conclude that PA homeostasis is essential for strawberry growth and development in response to environmental cues through the CSN5A-PAO5 model-mediated PA and H_2_O_2_ regulation.

### FvCSN5-mediated FvPAO5 degradation coordinates PA and ROS homeostasis to fine-tune strawberry growth

In plants, ROS functions as a double-edged sword, highlighting the importance of ROS homeostasis to accommodate diverse cellular roles and environmental responses (Farooq et al., 2019). Based on the available data and our present findings related to PAs, we summarize several key points, including (1) the dual roles of PAs as both ROS scavengers and triggers; (2) the multiple PA components with common and specialized cellular and biological processes, such as Put in photosynthesis, Spm in antioxidants, and Spd in both photosynthetic protection and oxidation resistance; (3) the essential role of Spd/Spm in reproductive growth and fitness; (4) the relationship between increased PAO5 activity and higher H_2_O_2_ levels; (5) the CSN5-mediated FvPAO5 degradation participates is plant vegetative and reproductive growth, suggesting that PAO5 may serve as a hub regulatory checkpoint for the balance between PAs and ROS, which are crucial for plant development and adaptation.

Earlier reports indicate that CSN5 is a key component of the COP9 signalosome and participates in various plant processes, including light signaling, circadian rhythm control, cell recycling, and IAA signaling (Chamovitz and Segal, 2001; Serino and Deng, 2003). CSN5 functions by the deneddylation of NEDD8 from the cullin subunit of CRL E3 ligases (Schwechheimer and Isono, 2010). This NEDD8-deneddylation is a post-translational modification that affects the activity of cullin-based and non-cullin substrates, while the CSN-mediated deneddylation is required but not essential for CRL-mediated processes (Mergner and Schwechheimer, 2014). Consistent with previous reports, the complete loss of CSN function results in seedling lethality (Stratmann and Gusmaroli, 2012). We observed that the inhibition of *FvCSN5A* expression led to developmental defects in strawberry leaves, flowers, and fruits (Fig. 3-5 and Fig. S4), and also affected PA levels and H_2_O_2_ production (Fig. 8-9). Interestingly, the overexpression of *FvPAO5* similarly affected plant development and pollen viability, akin to the phenotype of *FvCSN5A* RNAi plants (Fig. 10), suggesting a functional link between FvCSN5A with FvPAO5. This association is confirmed by a transient fruit transgenic experiment, showing that inhibiting *FaPAO5* expression could partially rescue the ripening phenotype of *FaCSN5A* RNAi fruits (Fig. 11). These data highlight that the post-translational modification of FvPAO5 by FvCSN5A contributes to the regulation of PA and ROS homeostasis, fine-tuning plant vegetative and reproductive growth.

The COP9 signalosome (CSN) controls the RUB/NEDD8 modification of several cullin-based E3 ligases, such as CUL□CDD□RBX1, and CUL□DDB1□COP1□SPA (Fischer et al., 2011; Qin et al., 2020; Dong et al., 2024). In the present study, we found that the expression pattern of *FvCUL1* was similar to that of *FvCSN5A* during fruit ripening, and the two proteins co-localized in the cytoplasm.

Specially, FvCSN5A interacted in vitro and in vivo with FvCUL1 (Fig. 2 and 7), a core component of E3 ubiquitin-protein ligase complexes. We further found the deneddylated and neddylated forms of FvCUL1 in the wild-type, with the protein levels of FvCUL1 markedly decreased in the *FvCSN5A* RNAi plants (Fig. 7). Additionally, FvCSN5A promoted the proteasome-dependent degradation of FvPAO5 in a ubiquitin-mediated manner (Fig. 6). Altogether, the CSN5A-CUL1-PAO5 pathway responsible for PA and H_2_O_2_ homeostasis, is crucial for strawberry plant growth and fruit ripening.

### An integrative model for FvCSN5A and FvPAO5 synergistically regulates the balance among Put, Spd and Spm essential to strawberry vegetative and reproductive growth by phytohormones and H_2_O_2_

To date, we have gained a more comprehensive understanding of the roles of PAs in embryo development, seed germination, seedling growth, reproductive development, and fruit ripening, as well as the relationship between PAs, hydrogen peroxide (H_2_O_2_), and phytohormone signaling. There is a close relationship between PAs and phytohormones in plant stomatal movement, survival fitness, and fruit ripening (Mo et al., 2020; Liu et al., 2023; Song et al., 2023; Gao et al., 2021, 2024a), with a particular emphasis on the interaction of PAs with ethylene in climacteric tomato fruit ripening and with ABA in non-climacteric strawberry fruit ripening (Gao et al., 2021). During the initiation of strawberry fruit ripening, increased ABA suppresses the expression of *FaPAO5* and reduces H_2_O_2_ production, ultimately facilitating the accumulation of Spm/ Spd and fruit ripening (Mo et al., 2020). Largely, PAs, especially Spm, regulate strawberry fruit ripening in an ABA-dominated, IAA-participating, and ethylene-coordinated manner (Guo et al., 2018). Similarly, in response to drought stress, the production of ABA controls PA homeostasis through both reducing Put biosynthesis and accelerating PA metabolism, in the process of which IAA is suggested to link to not only Put-regulating photosynthesis and oxidative phosphorylation, but also ABA-regulating sugar and flavonoid metabolism (Gao et al., 2024b). Thus, we consider that PAs, ABA, and IAA weave a core signaling network essential for plant growth and adaptation, focusing on the role of Put in photosynthesis and primary metabolism related to vegetative growth and plant fitness, and of Spd/Spm in secondary metabolism and survival fitness related to reproductive growth and fruit ripening. To a large extent, this may highlight PA metabolism acting as a feedback control mechanism in response to developmental and environmental stimuli.

In the present study, we found that the levels of IAA, ABA, and JA were suppressed in *FvCSN5A* RNAi plants, especially IAA with an 83% reduction; while the levels of both H_2_O_2_ production and *FvPAO5* expression were significantly elevated compared to the WT (Fig. 8 and 9). These data suggest that the inhibition of *FvCSN5A* expression significantly suppresses PA and IAA accumulation, accompanied by H_2_O_2_ burst, ultimately resulting in growth defects, retarded fruit ripening, and poor fertility phenotype. This is consistent with a previous report that OsPAO5 serves as a negative regulator for the mesocotyl elongation of rice seedlings, thus its knockout mutants facilitate faster seedling growth, ultimately increasing grain weight and yield by the higher accumulation of Spm and lower accumulation of H_2_O_2_ and JA (Lv et al., 2021). Also, in Arabidopsis, the loss of AtPAO5 function inhibits the CK-induced shoot meristem formation from lateral root primordia (Kaszler et al., 2023), potentially as a result of the interference of CK and IAA signaling in xylem differentiation (Alabdallah et al., 2017). Overall, the synergistic interaction between PAs and phytohormones lays a basis for the CSN5-PAO5 module responsible for the trade-off vegetative and reproductive growth mediated by H_2_O_2_.

Based on the available data and the present study, we propose a comprehensive model for the FvCSN5A-FvPAO5 coordinate regulation of PA metabolism and H_2_O_2_ homeostasis, which aligns with plant growth in response to environmental cues in strawberries (Fig. 12). The FvCSN5A-FvCUL1-FvPAO5 model provides novel insights into the integration of the COP9 signalosome into PA metabolism linked to IAA/ABA/JA by ROS, at least in model plant strawberries. Anyway, this may open a promising door for improvement of crop yield and quality through manipulating CSN5-PAO5 pair. Next, how the proposed model integrates early environmental cues by H_2_O_2_ underlying plant development and adaptation, is an intriguing question for further investigation.

**Figure 12.**
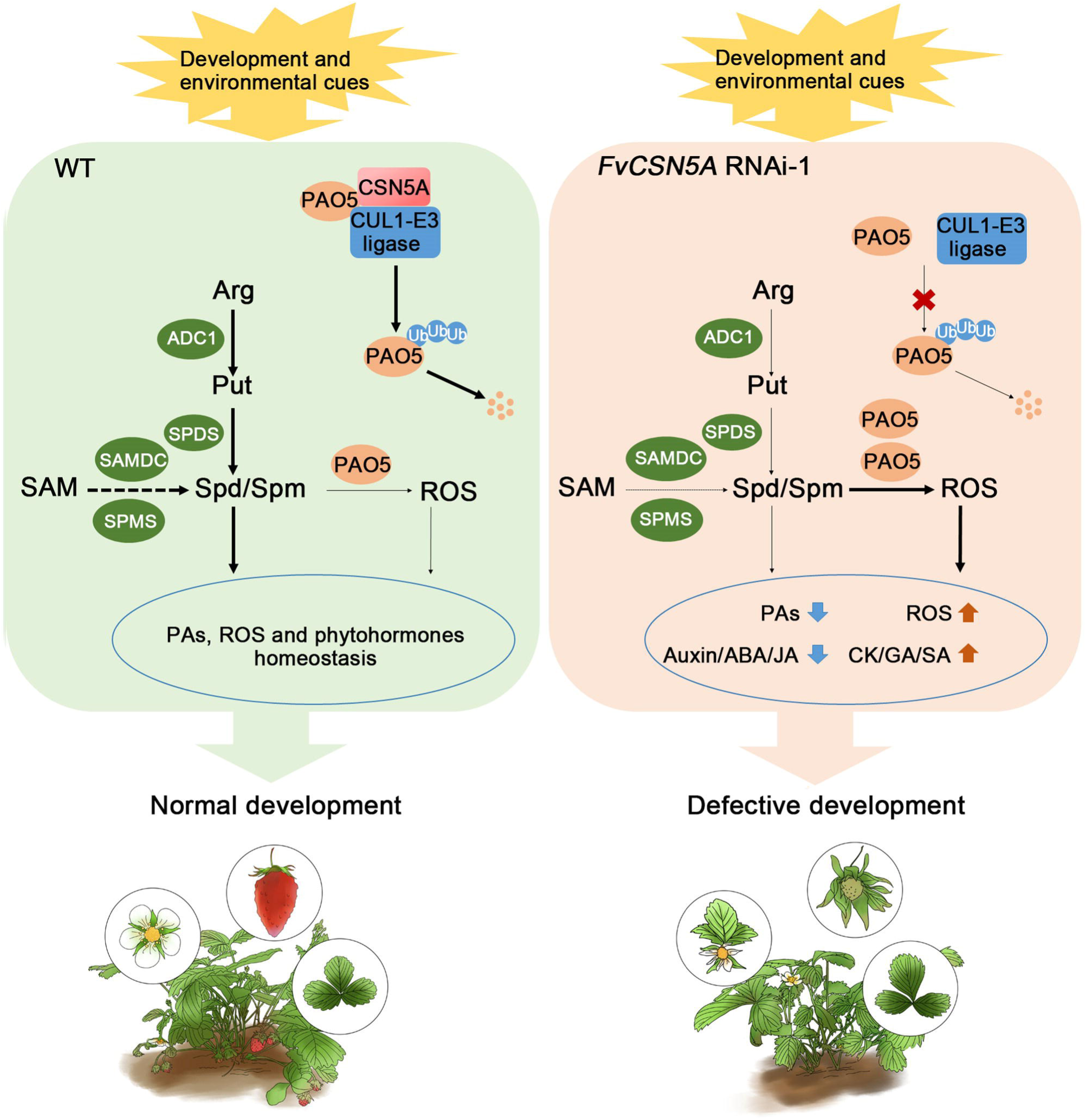
Proposed working model An equilibrium regulation for the composition and content of Put, Spd and Spm, is programmed by PA biosynthesis and degradation: Put-related key enzymes is ADC1; Spd/Spm-related key enzymes include SAMDC1, SPDS, and SPDM for biosynthesis and PAO5 for degradation. In response to environments, the two equilibria from Put/Spd/Spm metabolism to vegetative/reproductive growth are fine-tuned at the transcriptional, translational, and post-translational levels to optimize plant growth and breeding. The interaction of CSN5A with CUL1 mediates the degradation of PAO5, thereby rapidly controlling PA balance and H_2_O_2_ production, which can be integrated with various phytohormones. Finally, PAs, ROS, and phytohormones synergistically regulate strawberry development and adaptation. CSN5A is essential for the deneddylation of CUL1, potentially affecting the protein stability of CUL1 and the activity of E3 ubiquitin ligase. In *FvCSN5A* RNAi plants, CUL1 protein levels markedly decrease, while PAO5 levels increase. These changes lead to significant decreases in PA, Auxin, ABA, JA content, and excessive accumulation of H_2_O_2._ The homeostasis disruption in *FvCSN5A* RNAi plants may be responsible for the growth defects, slow fruit ripening and poor fertility phenotype.

## Materials and methods

### Plant materials and growth conditions

Octoploid strawberries (*Fragaria* × *ananassa* Duch. cv Benihoppe) grown in a greenhouse under natural sunlight conditions were used for transient transgenic experiments. Diploid strawberries (*Fragaria vesca*, Ruegen) and *Nicotiana benthamiana* were cultured in an artificial climate chamber at 25°C with a 16-h light/8-h dark photoperiod, 55% relative humidity, and a light intensity of 100 μmol m^-2^s^-1^.

### RNA isolation and RT-qPCR

Total RNA was extracted using the Plant Total RNA Mini-Extraction Kit (Magen, China) and qPCR Kit (YEASEN, China). RT-qPCR was performed on a Bio-Rad CFX96 system using the Trans Start Top Green qPCR Super Mix Kit (YEASEN, China) with *Actin* as an internal control.

### Yeast two-hybrid assay

For the yeast two-hybrid cDNA library screening, the coding sequence (CDS) of *FvPAO5* was cloned into a pGBKT7 vector. FvPAO5-BD was transformed into a yeast gold strain. The transformed yeast strain expressing *FvPAO5* was subsequently mated with a strawberry fruit cDNA library. After 22 to 24 h, the culture was plated onto SD/-Ade-His-Leu-Trp medium and incubated at 30°C for 3 to 5 days. Yeast colonies that grew on the medium were subsequently identified by PCR amplification and sequencing.

For the one-to-one yeast two-hybrid assay, the CDS of *FvPAO5* and *FvCUL1* were individually cloned into the pGBKT7 vector, while the CDS of *FvCSN5A* was cloned into the pGADT7 vector. BD-FvPAO5 and AD-FvCSN5A, as well as BD-FvCUL1 and AD-FvCSN5A, were transformed into yeast cells, respectively, as previously reported (Huang et al., 2024). The transformed yeast cells were cultured on SD/-Leu-Trp medium and incubated at 28°C for 3 days. Subsequently, the yeast cells were transferred to both SD/-Leu-Trp and SD/-Ade-His-Leu-Trp media, and incubated at 28°C for an additional 3 days before photos were taken. Autoactivation assays were performed for each bait and prey construct with the corresponding empty vectors to exclude potential false positives.

### Subcellular localization

The CDS regions of *FvCSN5A*, *FvCUL1*, and *FvPAO5* were constructed into the pCAMBIA1300-GFP vector and transiently expressed in tobacco leaves via Agrobacterium-mediated transformation. Nuclei were stained with DAPI in *N. benthamiana* leaves, and px-RK/CD3-983/CD3-983 was used as a marker for the peroxisome (Nelson et al., 2007). Visualization and imaging were performed using a Leica Stellaris 5 laser confocal microscope.

### Firefly luciferase complementation (FLC) assay

The CDS of *FvPAO5* and *FvCUL1* were cloned into pCAMBIA1300-NLUC, while the CDS of *FvCSN5A* was introduced into pCAMBIA1300-CLUC. These constructs were co-transfected into *Nicotiana benthamiana* leaves, and luminescence signals were recorded with a cold CCD camera 3 days after post-transfection.

### Bimolecular fluorescence complementation (BiFC) assay

The CDS of *FvCSN5A* was cloned into pSPYNE173, and the CDS of *FvPAO5* was cloned into pSPYCE. FvPAO5-YCE and FvCSN5A-YNE were co-transfected into *Nicotiana benthamiana* leaves. GFP signals were detected using laser confocal microscopy (Leica Stellaris 5) after 3 days.

### Co-Immunoprecipitation (CoIP) assay

Super1300:FvCSN5A-MYC and Super1300:FvPAO5-GFP constructs were cotransfected into *Nicotiana benthamiana* leaves. Leaves were collected after 2 days, and total proteins were extracted with a protein lysis buffer (100 mM Tris-HCl, pH 7.5, 50 mM NaCl, 5 mM EDTA, 1% TritonX-100, 1% Tween-20 and 1% NP-40). The extracted proteins were incubated with anti-MYC beads (AlpalifeBio, Cat. No.:KTSM1306) for 3 h. The target proteins were detected with an anti-MYC antibody (TransGen Biotech, Cat. No.:HT101-01; dilution 1:3000) and an anti-GFP antibody (TransGen Biotech, Cat. No.: HT801-01; dilution 1:3000).

### Co-immunoprecipitation mass spectrometry (MS)

This experiment was conducted according to the instructions of the ProteinA/G Immunoprecipitation Kit (Solarbio, Cat. No.: M2400). Total proteins were extracted from WT and *FvCSN5A* RNAi-1 leaves (10 g) using a protein extraction solution (100 mM HEPES, 1 mM EDTA, 10% glycerol, 1% TritonX-100, 1 mM DTT, 1 mM PMSF, 1% protease inhibitor cocktail, pH 7.8). Anti-FvPAO5 antibody was added to the pre-treated magnetic beads, and then total proteins were incubated with the beads for 1 h. The magnetic beads with bound proteins were washed 3 times using the extraction buffer. Finally, the magnetic beads with proteins were frozen in liquid nitrogen and used for mass spectrometry detection.

### GST pull-down assay

The CDS of *FvPAO5* was cloned into pGEX4T-1 (GST-tagged) and the CDS of *FvCSN5A* was cloned into pET-30a (His-tagged). Proteins were expressed in BL21 *Escherichia coli*. Recombinant FvPAO5-GST was purified and incubated with 20 µL glutathione Sepharose 4B in pull-down buffer (20 mM Tris–HCl, pH 7.5, 150 mM NaCl, 1 mM β-mercaptoethanol, 3 mM EDTA, pH 8.0, and 1% NP-40) for 1 h, and then FvCSN5A-His was added and incubated for 1 h. The proteins were then detected by immunoblotting using anti-His (TransGen Biotech, Cat. No.: HT501-01; dilution 1:3000) and anti-GST (TransGen Biotech, Cat. No.: HT601-01; dilution 1:3000) antibodies.

### Cell-free protein degradation assay

Cell-free degradation assays were carried out as described previously with little modification (Kong et al., 2015). The leaves of three-month-old ‘Ruegen’ strawberry seedlings were harvested and ground in liquid nitrogen. Total plant proteins were extracted using 0.5 g powder with 1.5 mL protein lysis buffer (100 mM HEPES, 1 mM EDTA, 10% glycerol, 1% TritonX-100, 1 mM DTT, 1 mM PMSF, 1% protease inhibitor cocktail, pH 7.8). The purified FvPAO5-GST protein (2.5 µg) and total strawberry protein were mixed with either DMSO or 50 µM MG132, respectively, and the samples were incubated at 20°C for various durations. Anti-GST (TransGen Biotech, Cat. No.: HT601-01; dilution 1:3000) and anti-Actin (BBI, Cat. No.: D110007-0100; dilution 1:3000) antibodies were used for immunoblotting detection.

### In vivo ubiquitination assay

In vivo ubiquitination assays were carried out as described previously (Ye et al., 2018). Three-month-old WT and *FvCSN5A* RNAi transgenic seedlings were cultured in liquid MS medium at 25°C for 5 h, with or without DMSO or 50 μM MG132 (stock solution prepared with 0.5% DMSO) and 100 μM CHX (cycloheximide, an inhibitor of protein synthesis). The total proteins were extracted as described above, and the total protein was incubated with ubiquitin-trap bead p62-agarose at 4°C for 4 h. Centrifuge at 1000 g for 1 min at 4 □, followed by discarding the supernatant, washing with 1 mL of PBS, and repeating this wash step twice. The ubiquitination signal was detected by an anti-PAO5 antibody (Beijing Protein Innovation; dilution 1:3000), while the input sample was detected by an anti-Actin antibody (BBI, Cat. No.: D110007-0100; dilution 1:3000).

### Protein changes in strawberry fruit development

The small green, white, and full red fruits of diploid strawberries were selected, achenes were removed, and samples were ground in liquid nitrogen. Total plant proteins were then extracted using 0.4 g powder with 200 µL protein lysis buffer (100 mM HEPES, 1 mM EDTA, 10% glycerol, 1% TritonX-100, 1 mM DTT, 1 mM PMSF, 1% protease inhibitor cocktail, pH 7.8). The target proteins were detected using the following antibodies: anti-FvPAO5 (Beijing Protein Innovation; dilution 1:3000), anti-FvCSN5A (Beijing Protein Innovation; dilution 1:3000), anti-CUL1 (PHYTO, Cat. No.: PHY1861S; dilution 1:3000), and anti-Actin (BBI, Cat. No.: D110007-0100; dilution 1:3000) antibody.

Transient transgene expression in strawberry fruits was performed as previously described (Chen et al., 2024). The CDS of *FaCSN5A* and *FaPAO5* was constructed into pCAMBIA1300-GFP vector to obtain the overexpression vector, while a 300 bp *FaCSN5A* and *FaPAO5* fragment was constructed into pK7GWIWG2 (II) vector for RNAi. Agrobacterium transformed *FaCSN5A* OE, control and *FaCSN5A* RNAi were incubated at 28°C with a final OD_600_ value of 0.5, and de-greening stage fruits were selected to inject. Ten fruits were used in each treatment. Photos were taken at 0, 2, 3, 4, and 5 days to record the phenotypes. Samples were harvested and achenes were removed 5 days after injection. The injection site was cut, and the injected parts were frozen in liquid nitrogen and stored at −80°C.

To explore the genetic interactions of strawberry CSN5A and PAO5, Agrobacterium transformed *FaCSN5A* RNAi, control, and *FaPAO5* RNAi were incubated at 28°C until reaching an OD_600_ value of 0.5, and de-greening stage fruits were selected injection according to control, *FaCSN5A* RNAi, *FaPAO5* RNAi, *FaCSN5A* RNAi and *FaPAO5* RNAi combinations. Each treatment included ten fruits. The fruit phenotype was recorded through photography at 0, 3, 6, and 7 days after injection. Samples were harvested and achenes were removed 7 days post-injection. The injection sites were excised, frozen in liquid nitrogen, and stored at −80°C.

### Stable transformation in diploid strawberry

The *FvCSN5A* RNAi (pK7GWIWG2 II: FvCSN5A_300 bp_-DsRed) and *FvPAO5* OE (Super1300:FvPAO5-GFP) constructs were transformed into the *Agrobacterium tumefaciens* strain GV3101, respectively. *Agrobacterium tumefaciens* cultures were incubated at 28°C, with a final OD_600_ value of 0.5, and then infiltrated into the diploid strawberry "Ruegen" using diploid strawberry leaf disks as described (Mao et al., 2022; Li et al., 2023; Lu et al., 2024). In this process, positive transgenic calli and regenerated plants were selected using DsRed or GFP fluorescence examined under a fluorescent protein observation lamp (LUYOR-3415RG). Fourteen *FvCSN5A* RNAi and ten *FvPAO5* OE transgenic seedlings were obtained. The experiment was carried out with T1 generation materials.

### In vitro germination of pollen grains

The anthers from freshly opened WT and *FvCSN5A* RNAi transgenic flowers were harvested. Pollen grains were released and incubated with pollen grain germination medium (10% sucrose, 0.002% boric acid, and 0.5% agar) at 22°C for 5 h. The pollen grains were taken to slides and photographed using a Leica DM750 microscope.

### Alexander’s staining

The anthers from freshly opened WT, *FvCSN5A* RNAi and *FvPAO5* OE transgenic flowers were harvested and Alexander’s staining solution (Coolabor, China) was applied to the anthers. The anthers were gently peeled off with tweezers, and the pollen grains were released into the staining solution for 6 h. Subsequently, the pollen grains were observed and photographed using a Leica DM750 microscope.

### Determination of anthocyanin contents

Samples (n=3) for determination of anthocyanin content was carried out according to the procedure described in the plant anthocyanin content detection kit (Solarbio, China). 0.1 g of strawberry sample powder was used in this experiment.

### Determination of soluble sugar contents

Soluble sugar content was determined by the previous method (Huang et al., 2019). Samples (n=3) of 0.5 g strawberry fruit powder was dissolved in 80% ethanol at 80°C for 30 min and repeated three times. After centrifugation, the supernatant was evaporated to dryness and dissolved in ultrapure water. The solution was filtered using LC-18 SPE and determined by HPLC. D-(+) glucose, D-(-) fructose and sucrose were used as standards.

### Determination of polyamines, 1, 3-diaminopropane (Dap), and phytohormones

PAs and Dap were determined according to the previous method (Guo et al., 2018) with little modification. Samples (n=3) of fresh strawberry leaves (0.5 g) were homogenized in 4 mL of 5% (v/v) cold perchloric acid and incubated at 4 °C for 1 h. Centrifugation was carried out at 4□ for 30 min at 12,000 g. The extracted supernatant was mixed with benzoyl chloride and subsequently analyzed for PAs and Dap using an HPLC system.

To determine the accumulation of Dap after exogenous Spd treatment, leaves were sampled 24 h post-treatment, and Dap content was carried out using the aforementioned methods. Each treatment was replicated three times.

Phytohormone contents were determined out according to the method described by Du et al. (2012) with minor modifications. Strawberry leaf samples (0.5 g FW) were ground into a fine powder in phytohormones and extracted with a modified Bieleski’s solvent (methanol/formic acid/water 15:1:4) overnight at 4°C. The extract was centrifuged at 12,000 rpm for 15 min at 4°C, followed by concentration and purification using primary secondary amine (PSA) and C_18_ tandem dual SPE cartridges. Finally, the extract was passed through a 0.45 µm organic filter membrane and analyzed by HPLC-MS/MS.

### DAB staining

For DAB (3,3’-Diaminobenzidine) staining, strawberry leaves, and anthers were incubated in DAB working staining solution (DAB color development kit, Solarbio, China) at room temperature for 4 h. Subsequently, the samples were decolorized using 95% ethanol at 80°C for 20 min. The DAB samples were then immersed in the preservation solution for 30 min. Pollen grains and leaves were photographed using a Leica DM750 microscope.

### Measurement of ROS production

For fluorescent detection of ROS using DCFH-DA, strawberry leaves and anthers were incubated with 5 µM DCFH-DA working staining solution (Solarbio, China) at room temperature for 10 min, and washed twice in PBS. Pollen grains and leaves were observed and photographed with a Leica Stellaris 5 laser confocal microscope.

Samples (n=3) for measurement of H_2_O_2_ following the method reported by Gay and Gebicki (2003), 0.2 g of strawberry leaf powder was incubated with pre-cooled acetone extraction buffer, and then extractant [(CCL4: CHCL3=3:1 (V:V)] and distilled water were added. The solution was centrifuged at 5000 r/min for 10 min, and the upper aqueous phase was taken to add the working reagent. After 30 min 30 □ water bath, the content of H_2_O_2_ was determined at A560 nm.

### Ploidy analysis

Cell nuclei were extracted and stained according to the guidelines of the CyStainR UV Precise P kit. The chromosome ploidy of young strawberry leaves from WT (diploid and tetraploid), *FvCSN5A* RNAi-1, *FvCSN5A* RNAi-2, Super1300-GFP (EV), Super1300:FvPAO5-GFP (*FvPAO5* OE-1 and *FvPAO5* OE-2) plants was analyzed using a CyFlowR space flow cytometer.

### Statistical analysis

All data are presented as means of at least three independent biological replicates. Statistical analyses were performed using SigmaPlot version 12.0 (Systat Software, USA). Data are presented as the analysis of variance (ANOVA) was performed. Asterisks indicate significant differences between treatments, assessed by Student’s t-test: *, P < 0.05; **, P < 0.01. Different letters indicate statistically significant differences at P < 0.05 as determined by ANOVA. A summary of the statistical analyses is provided in Supplemental Data Set 3.

## Author contributions

GF and GJX conceived the study and managed the projects. HY, GF, GJX, and SYY designed and guided the experiments. GJH, HY, and JGM performed most experiments. LWJ performed the determination of polyamines, Dap and H_2_O_2_. WQH performed yeast two-hybrid cDNA library screening assays. WJX assisted with the ploidy analysis. All authors analyzed the data. HY, GF, GJX, and GJH wrote and revised the article.

## Accession Numbers

The strawberry gene sequences from the *F. vesca* genome ver4.0 can be downloaded from the Genome Database for Rosaceae (http://ww.rosaceae.org) with accession numbers: FvPAO5, FvH4_7g01430; FvCSN5A, FvH4_2g36890; FvCUL1, FvH4_2g25090; FvCSN1, FvH4_7g1333; FvCSN2, FvH4_4g27570; FvCSN3, FvH4_6g52990; FvCSN4, FvH4_2g40000; FvCSN6A, FvH4_3g41360; FvCSN7, FvH4_7g10110; FvCSN7L, FvH4_5g19860; FvCSN8, FvH4_5g29770; FvSUT1, FvH4_5g33660; FvCHS, FvH4_7g01160; FvANS, FvH4_5g01170; FvPG, FvH4_7g01140; FvCEL, FvH4_4g33940; FvSS, FvH4_1g09360; FvADC1, FvH4_2g10570; FvSAMDC1, FvH4_6g45760; FvSPDS, FvH4_4g34680; FvSPMS, FvH4_7g31820.

## Supporting information

Supplemental Data Set1 Identification of FvPAO5 associated proteins by CO-IP-MS

Supplemental Data Set2 Primers used in this study

Supplemental Data Set3 Statistical analyses

## Data availability

The data underlying this article are available in the article and its online supplementary material.

## Acknowledgments

This work was supported by the National Natural Science Foundation of China (32030100; 32272648; 32072516; 32476225; 32372672) and R&D Program of Beijing Municipal Education Commission (KM202310020013). Partly supported by the open funds of the State Key Laboratory of Plant Environmental Resilience (SKLPERKF2407).

## Supplemental files

**Supplemental Figure 1.** F*v*CSN5A RNAi transgenic plants screening.

(**A)** The red calluses were *FvCSN5A* RNAi successful transgenic calluses, which were named *FvCSN5A* RNAi-1 and *FvCSN5A* RNAi-2.

(**B)** Visualization of DsRed accumulation in *FvCSN5A* RNAi transgenic plants (right). Bars, 1 cm.

**Supplemental Figure 2.** Peak plot of chromosome ploidy of *FvCSN5A* RNAi strawberries.

The young leaves of WT and *FvCSN5A* RNAi transgenic plants were used for chromosome ploidy analysis. The x-axes show the DNA contents, and the y-axes indicate cell numbers.

**Supplemental Figure 3.** Phenotype of *FvCSN5A* RNAi transgenic plants

(**A)** Leaf shape index of WT and *FvCSN5A* RNAi transgenic plants.

(**B)** Numbers of the serrations in WT and *FvCSN5A* RNAi transgenic plants.

(**C-D)** The images of pistil **(C)** and anther **(D)** in WT and *FvCSN5A* RNAi transgenic plants. Bars, 0.5 mm.

(**E-F)** Numbers of pistil **(E)** and anther **(F)** in WT and *FvCSN5A* RNAi transgenic plants.

(**G)** The expression of COP9 signalosome subunits in WT and *FvCSN5A* RNAi-1 fruits. Statistical significance of one-way ANOVA: *, P < 0.05; **, P < 0.01. Non-significant: ns

**Supplemental Figure 4.** Fruit Phenotype of *FvCSN5A* RNAi-2 transgenic fruits

(**A)** Reciprocal crosses between WT and *FvCSN5A* RNAi-2 transgenic fruits.

(**B)** Observation of fruit development process of two representative *FvCSN5A* RNAi-2 transgenic fruits. Photos were taken at 15, 17, 19, 21, 23, and 25 days after pollination (DAP) respectively to record the fruit phenotype. Bars, 0.5 cm.

**Supplemental Figure 5.** Manipulation of *FaCSN5A* expression affected physiological parameters and expression levels of ripening□related genes

(**A)** Phenotypes of *FaCSN5A* OE and *FaCSN5A* RNAi strawberry fruits. Agrobacterium GV3101 strains containing RNAi or overexpression *FaCSN5A* recombinant plasmids were injected into DG fruits attached to the plant. The fruit phenotype was recorded at 0, 2, 3, 4, and 5 days after injection. Bars, 1 cm.

(**B)** *FaCSN5A* expression levels in *FaCSN5A* OE and *FaCSN5A* RNAi fruits compared with the control.

(**C-E)** The anthocyanin content **(C)**, soluble sugar content **(D)**, and firmness **(E)** in *FaCSN5A* OE and *FaCSN5A* RNAi strawberry fruits.

(**F)** RT-qPCR was used to analyze the genes’ expression in *FaCSN5A* OE and *FaCSN5A* RNAi fruits. Relative expression levels were calculated using the 2^-△△Ct^ method. FaPG: polygalacturonase; FaCEL: cellulose; FaCHS: chalcone synthase; FaANS: anthocyanidin synthase; FaSUT1: sucrose transporter1; FaSS: sucrose synthase. FaADC1: arginine decarboxylase1; FaSAMDC1: S-adenosyl-methionine decarboxylase1; FaSPDS1: spermidine synthase1; FaSPMS: spermine synthase; FaPAO5: polyamine oxidase5.

Statistical significance of one-way ANOVA: *, P < 0.05; **, P < 0.01.

**Supplemental Figure 6.** Subcellular localization of FvCUL1

Subcellular localization of FvCUL1-GFP fusions in transiently transformed *N. benthamiana* leaves. All experiments were performed 48 h post-infiltration. Bars, 20 μm

**Supplemental Figure 7.** DAB dye of roots from WT and *FvCSN5A* RNAi transgenic plants. Seeds of WT and *FvCSN5A* RNAi transgenic plants were grown on MS medium for 3 months, and then the roots were harvested and stained with DAB.

**Supplemental Figure 8.** F*v*PAO5 OE transgenic plants screening and flower phenotype.

(**A)** Detection of *FvPAO5* OE (Super1300: FvPAO5-GFP) successful transgenic callus.

(**B)** Visualization of GFP fluorescence in *FvPAO5* OE plants. Bars, 1 cm.

(**C)** Peak plot of chromosome ploidy of *FvPAO5* OE strawberries.

(**D)** The flower phenotype of *FvPAO5* OE transgenic plants. Bars, 1cm.

**Supplemental Data Set 1** Identification of FvPAO5 associated proteins by CoIP/MS.

**Supplemental Data Set 2** The list of primers used in the experiments.

**Supplemental Data Set 3** Statistical analyses.

## References

Alabdallah O, Ahou A, Mancuso N, Pompili V, Macone A, Pashkoulov D, Stano P, Cona A, Angelini R, Tavladoraki P. The Arabidopsis polyamine oxidase/dehydrogenase 5 interferes with cytokinin and auxin signaling pathways to control xylem differentiation. Journal of Experimental Botany, 2017, 68(5): 997–1012.

Aloisi I, Piccini C, Cai G, Del Duca S. Male Fertility under Environmental Stress: Do Polyamines Act as Pollen Tube Growth Protectants?. International Journal of Molecular Sciences, 2022, 23(3): 1874.

Angelini R, Cona A, Federico R, Fincato P, Tavladoraki P, Tisi A. Plant amine oxidases "on the move": an update. Plant Physiology and Biochemistry, 2010, 48(7): 560–564.

Barry M, Früh K. Viral modulators of cullin RING ubiquitin ligases: culling the host defense. Science’s STKE, 2006, 2006(335): pe21.

Blázquez MA. Polyamines: Their Role in Plant Development and Stress. Annual Review of Plant Biology, 2024, 75. Available from: 10.1146/annurev-arplant-070623-110056

Chamovitz DA, Segal D. JAB1/CSN5 and the COP9 signalosome. A complex situation. EMBO Reports, 2001, 2(2): 96–101.

Chen X, Gao J, Shen Y. Abscisic acid controls sugar accumulation essential to strawberry fruit ripening via the FaRIPK1-FaTCP7-FaSTP13/FaSPT module. The Plant Journal, 2024, 119(3): 1400–1417.

Cope GA, Deshaies RJ. COP9 signalosome: A multifunctional regulator of SCF and other cullinDbased ubiquitin ligases. Cell, 2003, 114: 663–671.

Dong J, Li Y, Cheng S, Li X, Wei N. COP9 signalosome-mediated deneddylation of CULLIN1 is necessary for SCF^EBF1^ assembly in *Arabidopsis thaliana*. Cell reports, 2024, 43(1): 113638.

Du F, Ruan G, Liu H. Analytical methods for tracing plant hormones. Analytical and Bioanalytical Chemistry, 2012, 403(1): 55–74.

Farooq MA, Niazi AK, Akhtar J, Saifullah, Farooq M, Souri Z, Karimi N, Rengel Z. Acquiring control: The evolution of ROS-Induced oxidative stress and redox signaling pathways in plant stress responses. Plant Physiology and Biochemistry. 2019, 141: 353–369.

Fischer ES, Scrima A, Böhm K, Matsumoto S, Lingaraju GM, Faty M, Yasuda T, Cavadini S, Wakasugi M, Hanaoka F, Iwai S, Gut H, Sugasawa K, Thomä NH. The molecular basis of CRL4DDB2/CSA ubiquitin ligase architecture, targeting, and activation. Cell, 2011, 147: 1030–1039.

Fortes AM, Agudelo-Romero P, Pimentel D, Alkan N. Transcriptional modulation of polyamine metabolism in fruit species under abiotic and biotic stress. Frontiers in Plant Science, 2019, 10: 816.

Furumoto T, Yamaoka S, Kohchi T, Motose H, Takahashi T. Thermospermine is an evolutionarily ancestral phytohormone required for organ development and stress responses in *Marchantia polymorpha*. Plant & Cell Physiology, 2024, 65(3): 460–471.

Gao F, Mei XR, Li YZ, Guo JX, Shen YY. Update on the roles of polyamines in fleshy fruit ripening, senescence, and quality. Frontiers in Plant Science, 2021, 12: 610313.

Gao F, Li JY, Li WJ, Shi S, Song SH, Shen YY, Guo JX. Abscisic acid and polyamines coordinately regulate strawberry drought responses. Plant Stress, 2024a, 11: 100387.

Gao F, Guo J, Shen Y. Advances from chlorophyll biosynthesis to photosynthetic adaptation, evolution and signaling. Plant Stress, 2024b, 12: 100470.

Gay CA, Gebicki JM. Measurement of protein and lipid hydroperoxides in biological systems by the ferric-xylenol orange method. Anal Biochemistry, 2003, 315(1):29–35.

Groppa MD, Benavides MP. Polyamines and abiotic stress: recent advances. Amino Acids, 2008, 34(1): 35–45.

Guo J, Wang S, Yu X, Dong R, Li Y, Mei X, Shen Y. Polyamines regulate strawberry fruit ripening by abscisic Acid, auxin, and ethylene. Plant Physiology. 2018, 177(1): 339–351.

Gupta K, Sengupta A, Chakraborty M, Gupta B. Hydrogen peroxide and polyamines act as double edged swords in plant abiotic stress responses. Frontiers in Plant Science, 2016, 7: 1343.

Gusmaroli G, Feng S, Deng XW. The Arabidopsis CSN5A and CSN5B subunits are present in distinct COP9 signalosome complexes, and mutations in their JAMM domains exhibit differential dominant negative effects on development. The Plant Cell, 2004, 16: 2984–3001.

Hotton SK, Callis J. Regulation of cullin RING ligases. Annual Review of Plant Biology, 2008, 59: 467–489.

Huang F, Sun M, Yao Z, Zhou J, Bai Q, Chen X, Huang Y, Shen Y. Protein kinase FaSnRK2.6 phosphorylates transcription factor FabHLH3 to regulate anthocyanin homeostasis during strawberry fruit ripening. Journal of Experimental Botany, 2024, 29: erae250.

Huang X, Hetfeld BK, Seifert U, Kahne T, Kloetzel PM, Naumann M, BechDOtschir D, Dubiel W. Consequences of COP9 signalosome and 26S proteasome interaction. The FEBS Journal, 2005, 272: 3909–3917.

Huang Y, Xu PH, Hou BZ, Shen YY. Strawberry tonoplast transporter, FaVPT1, controls phosphate accumulation and fruit quality. Plant Cell & Environment, 2019, 42(9): 2715–2729.

Jasso-Robles FI, Gonzalez ME, Pieckenstain FL, Ramírez-García JM, Guerrero-González ML, Jiménez-Bremont JF, Rodríguez-Kessler M. Decrease of Arabidopsis PAO activity entails increased RBOH activity, ROS content and altered responses to *Pseudomonas*. Plant Science, 2020, 292: 110372.

Jin D, Li B, Deng XW, Wei N. Plant COP9 signalosome subunit 5, CSN5. Plant Science, 2014, 224: 54–61.

Kaszler N, Benkő P, Molnár Á, Zámbori A, Fehér A, Gémes K. Absence of Arabidopsis polyamine oxidase 5 influences the cytokinin-induced shoot meristem formation from lateral root primordia. Plants, 2023, 12(3): 454.

Kong L, Cheng J, Zhu Y, Ding Y, Meng J, Chen Z, Xie Q, Guo Y, Li J, Yang S, Gong Z. Degradation of the ABA co-receptor ABI1 by PUB12/13 U-box E3 ligases. Nature Communications, 2015, 6: 8630.

Kwok SF, Solano R, Tsuge T, Chamovitz DA, Ecker JR, Matsui M, Deng XW. Arabidopsis homologs of a cDJun coactivator are present both in monomeric form and in the COP9 complex, and their abundance is differentially affected by the pleiotropic cop/det/fus mutations. The Plant Cell, 1998, 10: 1779–1790.

Li W, Zhang J, Sun H, Wang S, Chen K, Liu Y, Li H, Ma Y, Zhang Z. FveRGA1, encoding a DELLA protein, negatively regulates runner production in *Fragaria vesca*. Planta, 2018, 247(4): 941–951.

Li X, Martín-Pizarro C, Zhou L, Hou B, Wang Y, Shen Y, Li B, Posé D, Qin G. Deciphering the regulatory network of the NAC transcription factor FvRIF, a key regulator of strawberry (*Fragaria vesca*) fruit ripening. The Plant Cell, 2023, 35(11): 4020–4045.

Liao X, Li M, Liu B, Yan M, Yu X, Zi H, Liu R, Yamamuro C. Interlinked regulatory loops of ABA catabolism and biosynthesis coordinate fruit growth and ripening in woodland strawberry. Proceedings of the National Academy of Sciences of the United States of America, 2018, 115(49): E11542–E11550.

Lingaraju GM, Bunker RD, Cavadini S, Hess D, Hassiepen U, Renatus M, Fischer ES, Thoma NH. Crystal structure of the human COP9 signalosome. Nature, 2014, 512: 161–165.

Liu J, Nada K, Pang X, Honda C, Kitashiba H, Moriguchi T. Role of polyamines in peach fruit development and storage. Tree Physiology, 2006, 26(6): 791–798.

Liu X, Reitsma JM, Mamrosh JL, Zhang Y, Straube R, Deshaies RJ. Cand1Dmediated adaptive exchange mechanism enables variation in FDbox protein expression. Molecular Cell, 2018, 69(5): 773–786.

Liu XD, Zeng YY, Zhang XY, Tian XQ, Hasan MM, Yao GQ, Fang XW. Polyamines inhibit abscisic acid-induced stomatal closure by scavenging hydrogen peroxide. Physiologia Plantarum, 2023, 175(2): e13903.

Lu J, Yu J, Liu P, Gu J, Chen Y, Zhang T, Li J, Wang T, Yang W, Lin R, Wang F, Qi M, Li T, Liu Y. Ubiquitin-mediated degradation of SlPsbS regulates low night temperature tolerance in tomatoes. Cell Reports, 2024, 43(10): 114757.

Lu R, Hu S, Feng J, Liu Z, Kang C. The AP2 transcription factor BARE RECEPTACLE regulates floral organogenesis via auxin pathways in woodland strawberry. The Plant Cell, 2024, koae270, 10.1093/plcell/koae270

Lv Y, Shao G, Jiao G, Sheng Z, Xie L, Hu S, Tang S, Wei X, Hu P. Targeted mutagenesis of POLYAMINE OXIDASE 5 that negatively regulates mesocotyl elongation enables the generation of direct-seeding rice with improved grain yield. Molecular plant, 2021, 14(2): 344–351.

Lyapina S, Cope G, Shevchenko A, Serino G, Tsuge T, Zhou C, Wolf DA, Wei N, Shevchenko A, Deshaies RJ. Promotion of NEDD-CUL1 conjugate cleavage by COP9 signalosome. Science, 2001, 292: 1382–1385.

Mao W, Han Y, Chen Y, Sun M, Feng Q, Li L, Liu L, Zhang K, Wei L, Han Z, Li B. Low temperature inhibits anthocyanin accumulation in strawberry fruit by activating FvMAPK3-induced phosphorylation of FvMYB10 and degradation of chalcone synthase 1. The Plant Cell, 2022, 34(4): 1226–1249.

Mayor-Ruiz C, Jaeger MG, Bauer S, Brand M, Sin C, Hanzl A, Mueller AC, Menche J, Winter GE. Plasticity of the cullin-RING ligase repertoire shapes sensitivity to ligand-induced protein degradation. Molecular Cell, 2019, 75: 849–858.

Mergner J, Schwechheimer C. The NEDD8 modification pathway in plants. Frontiers in Plant Science. 2014, 5:103.

Mo AW, Xu T, Bai Q, Shen YY, Gao F, Guo JX. FaPAO5 regulates Spm/Spd levels as a signaling during strawberry fruit ripening. Plant Direct, 2020, 4(5): e00217.

Nandy S, Das T, Tudu CK, Mishra T, Ghorai M, Gadekar VS, Anand U, Kumar M, Behl T, Shaikh NK, Jha NK, Shekhawat MS, Pandey DK, Dwivedi P, Radha, Dey A. Unravelling the multi-faceted regulatory role of polyamines in plant biotechnology, transgenics and secondary metabolomics. Applied Microbiology and Biotechnology, 2022, 106(3): 905–929.

Napieraj N, Janicka M, Reda M. Interactions of polyamines and phytohormones in plant response to abiotic stress. Plants, 2023, 12(5): 1159.

Navakoudis E, Kotzabasis K. Polyamines: Α bioenergetic smart switch for plant protection and development. Journal of Plant Physiology, 2022, 270: 153618.

Nelson BK, Cai X, Nebenführ A. A multicolored set of in vivo organelle markers for co-localization studies in Arabidopsis and other plants. The Plant Journal, 2007, 51(6): 1126–1136.

Paschalidis KA, Roubelakis-Angelakis KA. Sites and regulation of polyamine catabolism in the tobacco plant. Correlations with cell division/expansion, cell cycle progression, and vascular development. Plant Physiology, 2005, 138(4): 2174–2184.

Pottosin I, Velarde-Buendía AM, Bose J, Zepeda-Jazo I, Shabala S, Dobrovinskaya O. Cross-talk between reactive oxygen species and polyamines in regulation of ion transport across the plasma membrane: implications for plant adaptive responses. Journal of Experimental Botany, 2014, 65(5): 1271–1283.

Qin N, Xu D, Li J, Deng XW. COP9 signalosome: Discovery, conservation, activity, and function. Journal of integrative plant biology, 2020, 62(1): 90–103.

Reitsma JM, Liu X, Reichermeier KM, Moradian A, Sweredoski MJ, Hess S, Deshaies RJ. Composition and regulation of the cellular repertoire of SCF ubiquitin ligases. Cell, 2017, 171(6): 1326–1339.

Russell DH. The roles of polyamines, putrescine, spermidine and spermine in normal and malignant tissues. Life Science, 1973, 13: 1635–1647.

Schwechheimer C, Serino G, Deng XW. Interactions of the COP9 signalosome with the E3 ubiquitin ligase SCFTIR1 in mediating auxin response. Science, 2001, 292: 1379–1382.

Schwechheimer C, Isono E. The COP9 signalosome and its role in plant development. European Journal of Cell Biology. 2010, 89(2): 157–162.

Serino G, Deng XW. The COP9 signalosome: regulating plant development through the control of proteolysis. Annual Review of Plant Biology, 2003, 54(1):165–182.

Shang Y, Wang K, Sun S, Zhou J, Yu JQ. COP9 Signalosome CSN4 and CSN5 subunits are involved in jasmonate-dependent defense against root-knot nematode in tomato. Frontier in Plant Science, 2019, 10: 1223.

Shen ZF, Li L, Wang JY, Liao J, Zhang YR, Zhu XM, Wang ZH, Lu JP, Liu XH, Lin FC. CSN5 inhibits autophagy by regulating the ubiquitination of Atg6 and Tor to mediate the pathogenicity of *Magnaporthe oryzae*. Cell Communication and Signaling, 2024, 22(1): 222.

Siddappa S, Marathe GK. What we know about plant arginases? Plant Physiology and Biochemistry, 2020, 156: 600–610.

Song J, Sun P, Kong W, Xie Z, Li C, Liu JH. SnRK2.4-mediated phosphorylation of ABF2 regulates ARGININE DECARBOXYLASE expression and putrescine accumulation under drought stress. New Phytologist, 2023, 238(1): 216–236.

Stratmann JW, Gusmaroli G. Many jobs for one good cop - the COP9 signalosome guard development and defense. Plant Science, 2012, 185-186: 50–64.

Verma R, Aravind L, Oania R, McDonald WH, Yates JR 3rd, Koonin EV, Deshaies RJ. Role of Rpn11 metalloprotease in deubiquitination and degradation by the 26S proteasome. Science. 2002, 298(5593): 611–615.

Wang W, Paschalidis K, Feng JC, Song J, Liu JH. Polyamine catabolism in plants: A universal process with diverse functions. Frontiers in Plant Science, 2019, 10: 561.

Wei N, Deng XW. The COP9 signalosome. Annual Review of Cell and Developmental Biology, 2003, 19(1): 261–286.

Ye Q, Wang H, Su T, Wu WH, Chen YF. The Ubiquitin E3 Ligase PRU1 Regulates WRKY6 Degradation to Modulate Phosphate Homeostasis in Response to Low-Pi Stress in Arabidopsis. The Plant Cell, 2018, 30(5):1062–1076.

Yu Z, Jia DY, Liu TB. Polyamine oxidases play various roles in plant development and abiotic stress tolerance. Plants, 2019, 8(6): 184.

Zhan ZN, Wang N, Chen ZM, Zhang YX, Geng KQ, Li DM, Wang ZP. Effects of water stress on endogenous hormones and free polyamines in different tissues of grapevines (*Vitis vinifera* L. cv. ‘Merlot’). Functional Plant Biology, 2023, 50(12): 993–1009.

Zhao JQ, Wang XF, Pan XB, Jiang QQ, Xi ZM. Exogenous putrescine alleviates drought stress by altering reactive oxygen species scavenging and biosynthesis of polyamines in the seedlings of Cabernet Sauvignon. Frontiers in Plant Science, 2021, 12: 767992.

Zhong M, Yue L, Liu W, Qin H, Lei B, Huang R, Yang X, Kang Y. Genome-wide identification and characterization of the polyamine uptake transporter (Put) gene family in tomatoes and the role of Put2 in response to salt stress. Antioxidants, 2023, 12(2): 228.

