## Supplemental Figures for "Strawberry COP9 signalosome FvCSN5A regulates plant development and fruit ripening by facilitating polyamine oxidase FvPAO5 degradation to control polyamine and H_2_O_2_ homeostasis"

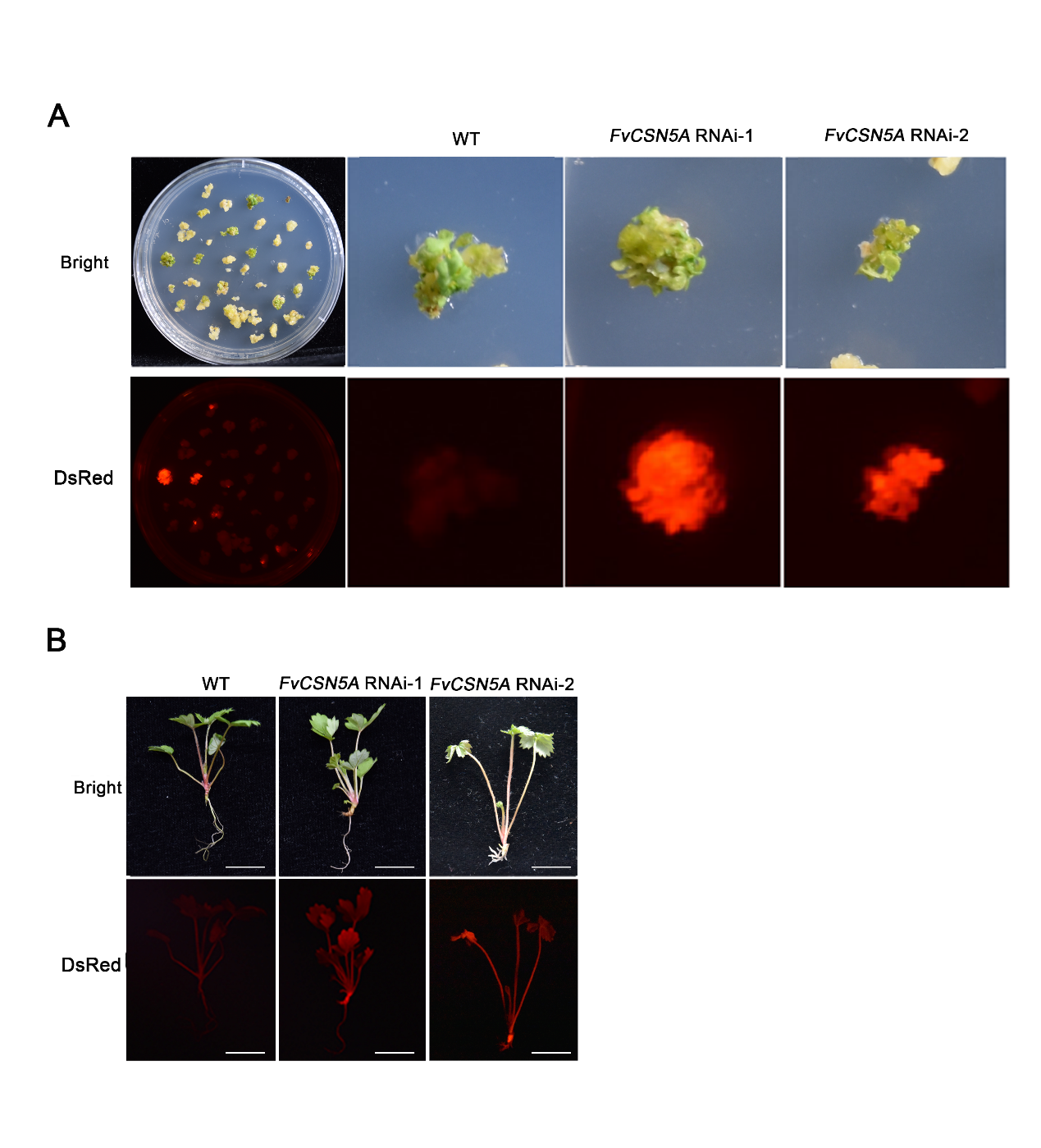

### Supplemental Figure 1. *FvCSN5A* RNAi transgenic plants screening.

### (A) The red calluses were *FvCSN5A* RNAi successful transgenic calluses, which were named *FvCSN5A* RNAi-1 and *FvCSN5A* RNAi-2.

**(B)** Visualization of DsRed accumulation in *FvCSN5A* RNAi transgenic plants (right). Bars, 1 cm.

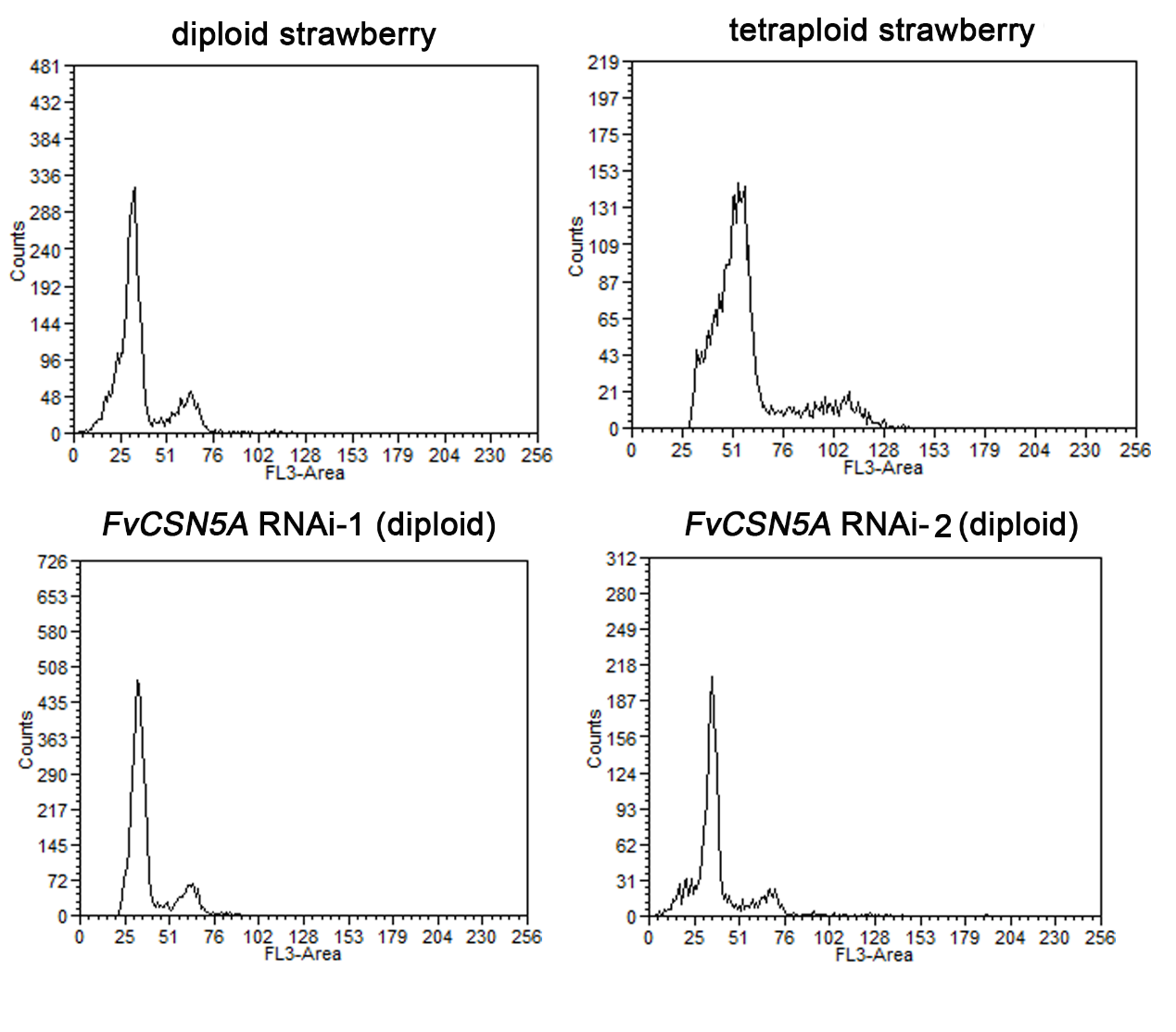

**Supplemental Figure 2.** Peak plot of chromosome ploidy of *FvCSN5A* RNAi strawberries.

The young leaves of WT and *FvCSN5A* RNAi transgenic plants were used for chromosome ploidy analysis. The x-axes show the DNA contents, and the y-axes indicate cell numbers.

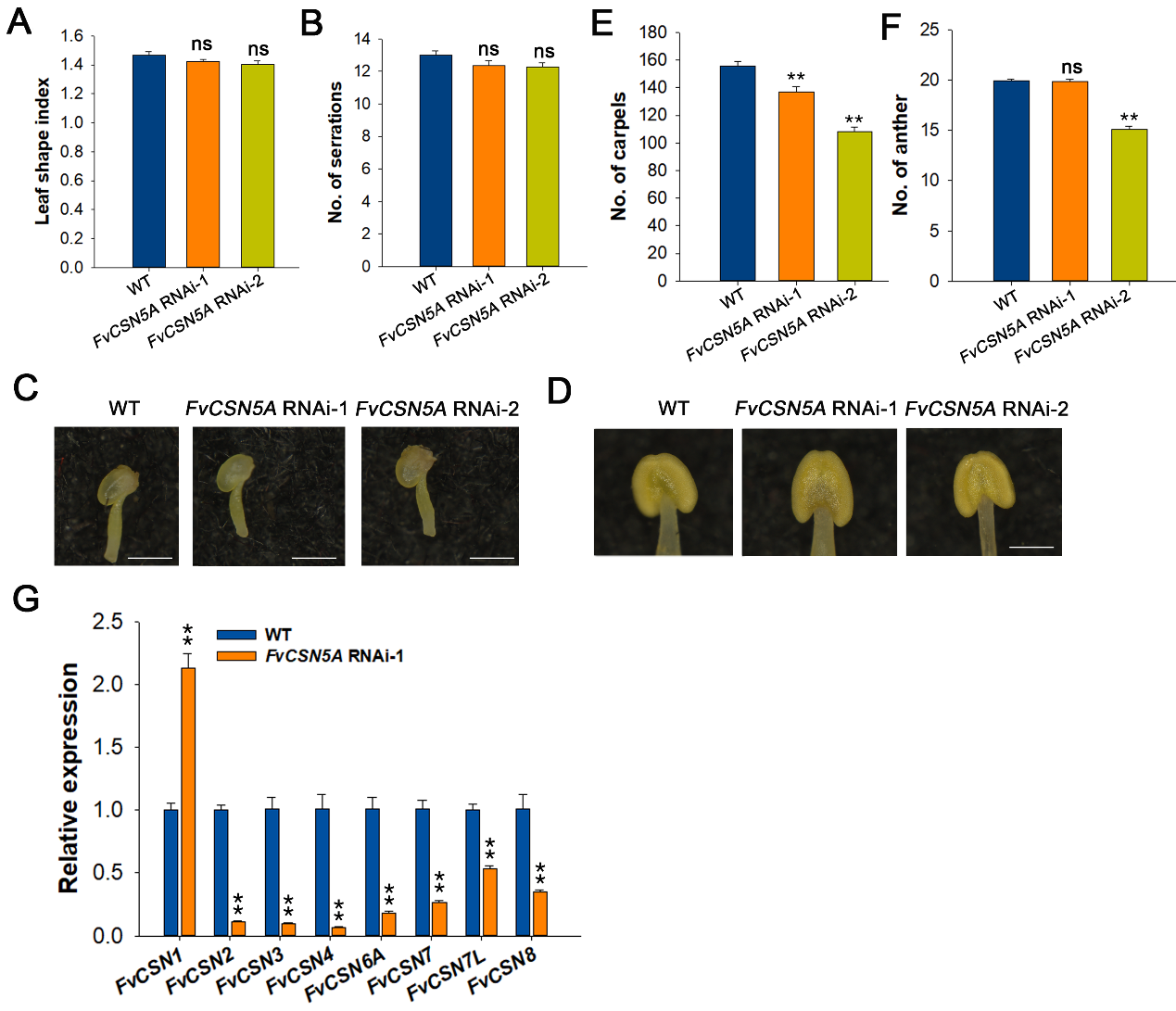

**Supplemental Figure 3.** Phenotype of *FvCSN5A* RNAi transgenic plants

**(A)** Leaf shape index of WT and *FvCSN5A* RNAi transgenic plants.

**(B)** Numbers of the serrations in WT and *FvCSN5A* RNAi transgenic plants.

**(C-D)** The images of pistil **(C)** and anther **(D)** in WT and *FvCSN5A* RNAi transgenic plants. Bars, 0.5 mm.
**(E-F)** Numbers of pistil **(E)** and anther **(F)** in WT and *FvCSN5A* RNAi transgenic plants.

**(G)** The expression of COP9 signalosome subunits in WT and *FvCSN5A* RNAi-1 fruits.

Statistical significance of one-way ANOVA: *, P < 0.05; **, P < 0.01. Non-significant: ns.

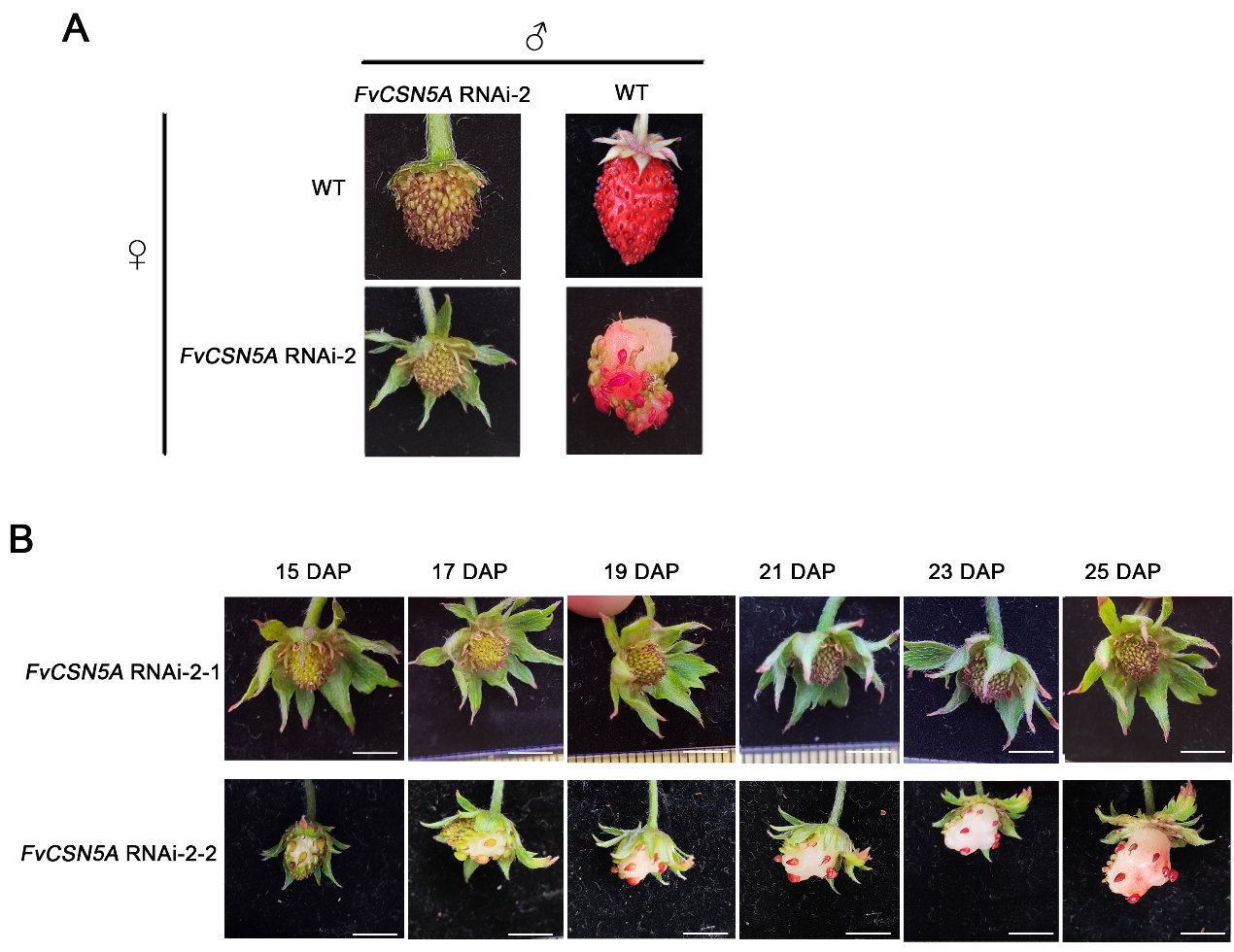

### Supplemental Figure 4. Fruit Phenotype of *FvCSN5A* RNAi-2 transgenic fruits

### (A) Reciprocal crosses between WT and *FvCSN5A* RNAi-2 transgenic fruits.

### (B) Observation of fruit development process of two representative *FvCSN5A* RNAi-2 transgenic fruits. Photos were taken at 15, 17, 19, 21, 23, and 25 days after pollination (DAP) respectively to record the fruit phenotype. Bars, 0.5 cm.

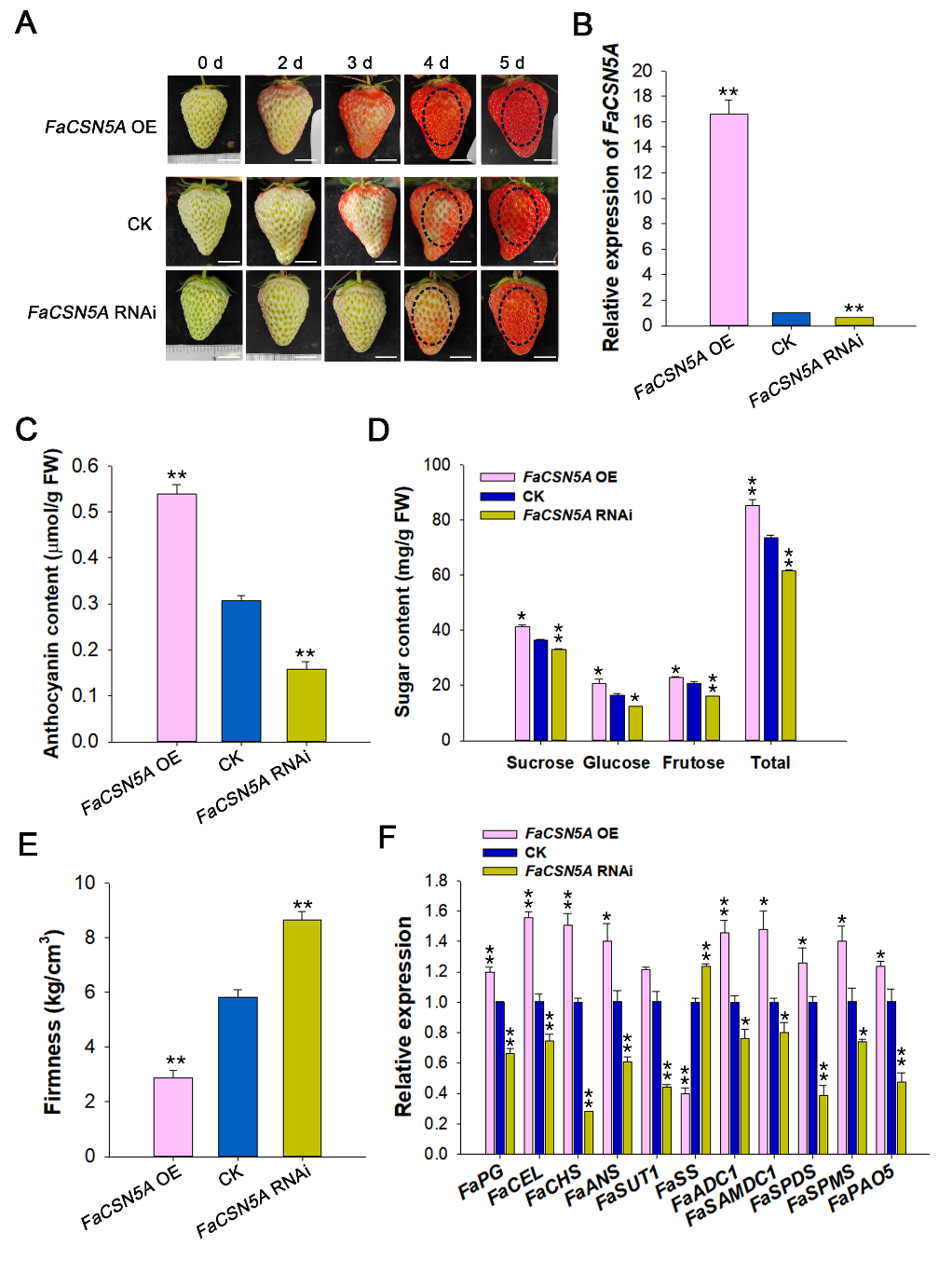

### Supplemental Figure 5. Manipulation of *FaCSN5A* expression affected physiological parameters and expression levels of ripening‐related genes

**(A)** Phenotypes of *FaCSN5A* OE and *FaCSN5A* RNAi strawberry fruits. Agrobacterium GV3101 strains containing RNAi or overexpression *FaCSN5A* recombinant plasmids were injected into DG fruits attached to the plant. The fruit phenotype was recorded at 0, 2, 3, 4, and 5 days after injection. Bars, 1 cm.

**(B)** *FaCSN5A* expression levels in *FaCSN5A* OE and *FaCSN5A* RNAi fruits compared with the control.

**(C-E)** The anthocyanin content **(C)**, soluble sugar content **(D)**, and firmness **(E)** in *FaCSN5A* OE and *FaCSN5A* RNAi strawberry fruits.

**(F)** RT-qPCR was used to analyze the genes’ expression in *FaCSN5A* OE and *FaCSN5A* RNAi fruits. Relative expression levels were calculated using the 2^-△△Ct^ method. FaPG: polygalacturonase; FaCEL: cellulose; FaCHS: chalcone synthase; FaANS: anthocyanidin synthase; FaSUT1: sucrose transporter1; FaSS: sucrose synthase. FaADC1: arginine decarboxylase1; FaSAMDC1: S-adenosyl-methionine decarboxylase1; FaSPDS1: spermidine synthase1; FaSPMS: spermine synthase; FaPAO5: polyamine oxidase5.

Statistical significance of one-way ANOVA: *, P < 0.05; **, P < 0.01.

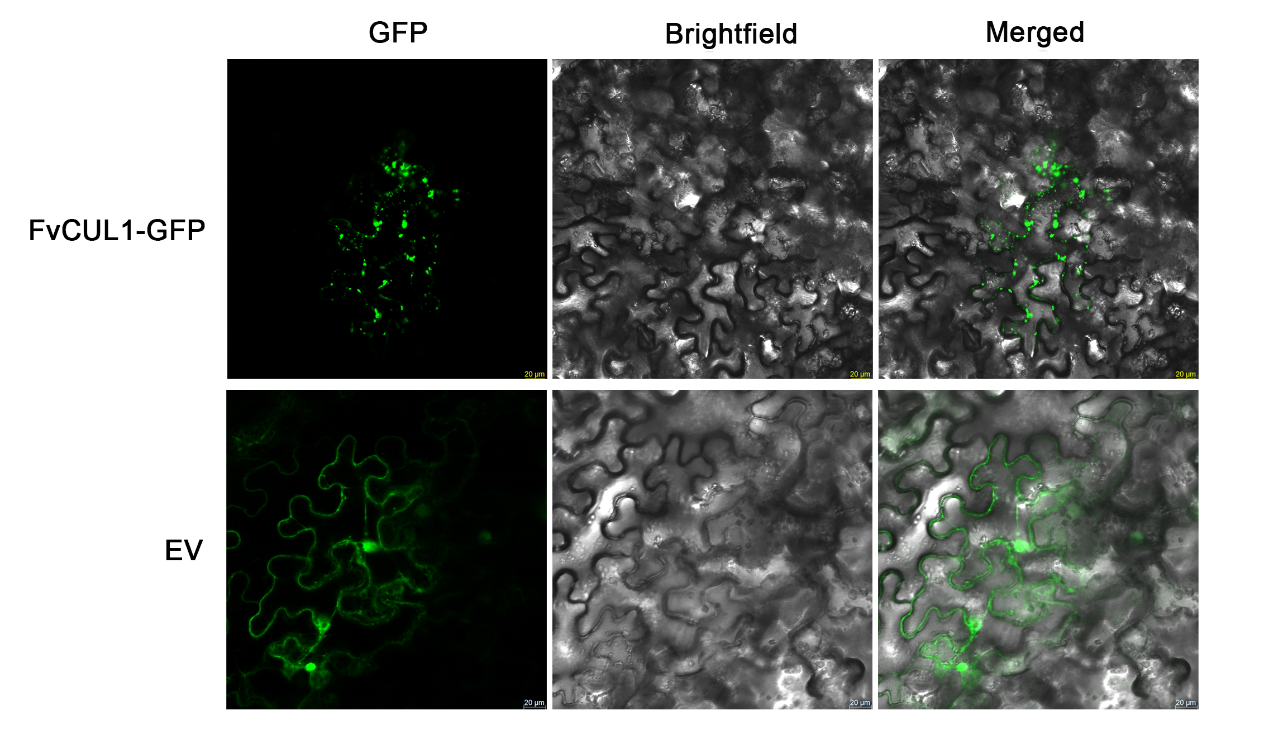

**Supplemental Figure 6.** Subcellular localization of FvCUL1

Subcellular localization of FvCUL1-GFP fusions in transiently transformed *N. benthamiana* leaves. All experiments were performed 48 h post-infiltration. Bars, 20 μm.

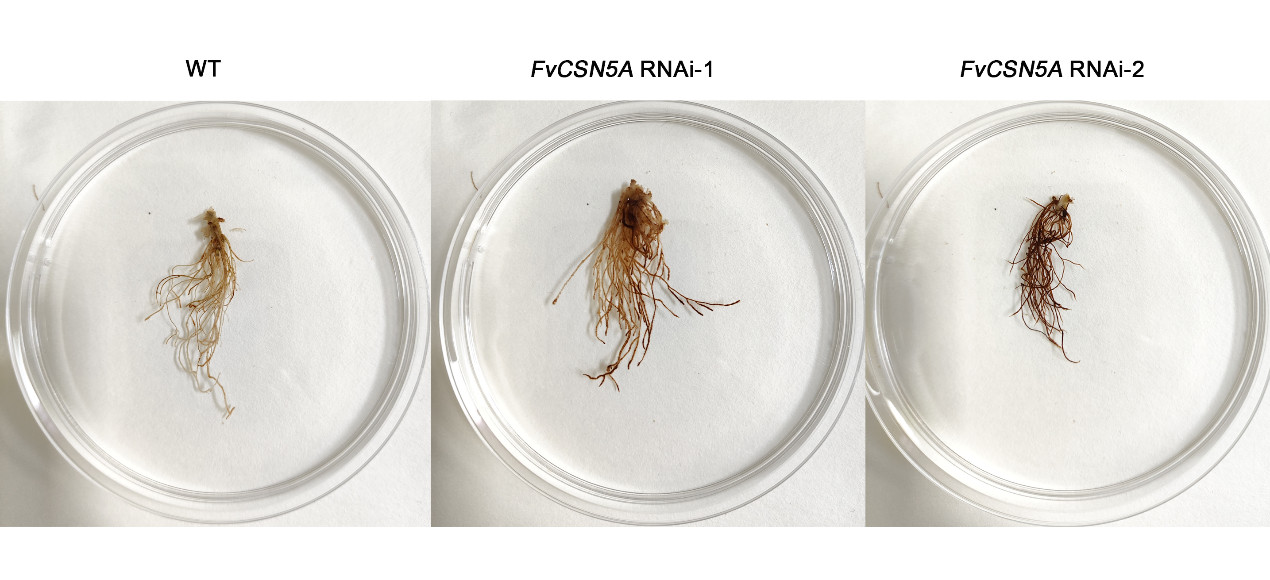

**Supplemental Figure 7.** DAB dye of roots from WT and *FvCSN5A* RNAi transgenic plants.

Seeds of WT and *FvCSN5A* RNAi transgenic plants were grown on MS medium for 3 months, and then the roots were harvested and stained with DAB.

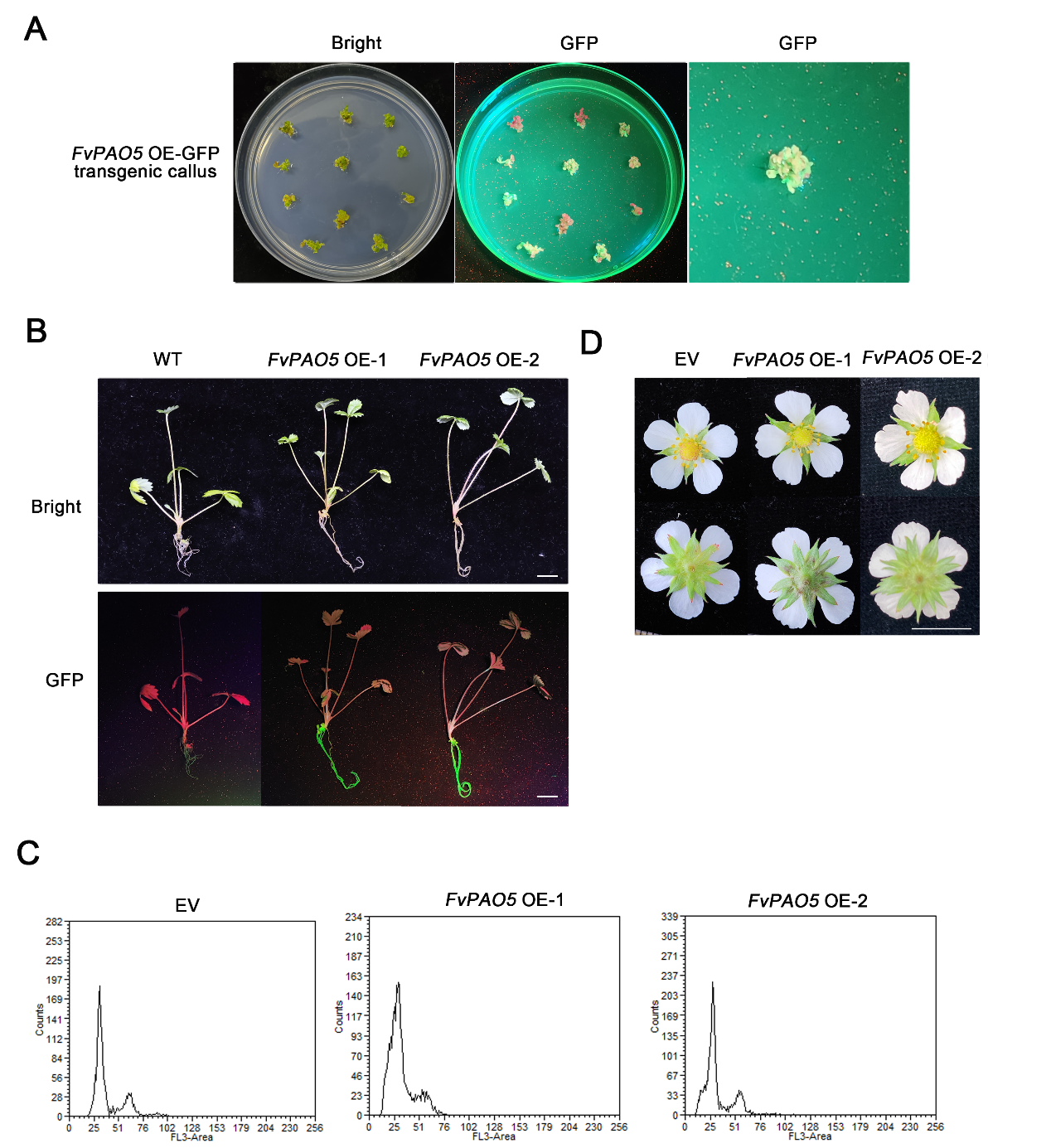

**Supplemental Figure 8.** *FvPAO5* OE transgenic plants screening and flower phenotype.

**(A)** Detection of *FvPAO5* OE (Super1300: FvPAO5-GFP) successful transgenic callus.

**(B)** Visualization of GFP fluorescence in *FvPAO5* OE plants. Bars, 1 cm.

**(C)** Peak plot of chromosome ploidy of *FvPAO5* OE strawberries.

**(D)** The flower phenotype of *FvPAO5* OE transgenic plants. Bars, 1cm.
